# Adsorption-driven deformation and landing-footprints of the RBD proteins in SARS-CoV-2 variants onto biological and inanimate surfaces

**DOI:** 10.1101/2024.01.15.575706

**Authors:** Antonio Bosch, Horacio V. Guzman, Rubén Pérez

**Affiliations:** Departamento de Física Téorica de la Materia Condensada, Universidad Autónoma de Madrid, E-28049 Madrid, Spain; Condensed Matter Physics Center (IFIMAC), Universidad Autónoma de Madrid, E-28049 Madrid, Spain; Department of Theoretical Physics, Jožef Stefan Institute, SI-1000 Ljubljana, Slovenia

**Author notes:** These authors share first authorship (Antonio Bosch, Horacio V. Guzman).

## Abstract

Respiratory viruses, carried through airborne microdroplets, frequently adhere to surfaces, including plastics and metals. However, our understanding of the interactions between viruses and materials remains limited, particularly in scenarios involving polarizable surfaces. Here, we investigate the role of receptor-binding domain (RBD) mutations on the adsorption of SARS-CoV-2 to hydrophobic and hydrophilic surfaces employing molecular simulations. To contextualize our findings, we contrast the interactions on inanimate surfaces with those on native-biological interfaces, specifically the ACE2 receptor. Notably, we identify a twofold increase in structural deformations for the protein’s receptor binding motif onto the inanimate surfaces, indicative of enhanced shock-absorbing mechanisms. Furthermore, the distribution of amino acids (landing-footprints) on the inanimate surface reveals a distinct regional asymmetry relative to the biological interface. In spite of the H-bonds formed at the hydrophilic substrate, the simulations consistently show a higher number of contacts and interfacial area with the hydrophobic surface, with the WT RBD adsorbed more strongly than the delta or omicron RBDs. In contrast, the adsorption of delta and omicron to hydrophilic surfaces was characterized by a distinctive hopping-pattern. The novel shock-absorbing mechanisms identified in the virus adsorption on inanimate surfaces could lead current experimental efforts in the design of virucidal surfaces.

## Introduction

Respiratory viruses are airborne and commonly form microdroplets that can be easily adsorbed onto substrates of different materials, namely, polymers, metals, textiles, and glasses, among other prophylactic materials. The recent pandemic has brought the SARS-CoV-2 virus to the spotlight, because of its higher transmission rates.^1–4^ The scientific community has delivered a rapid response in several research fields, from obtaining high-resolution structures of the virus, ^1,2,5^ to developing vaccines and therapies,^6–8^ passing by several endeavours to elucidate the behavior and weaknesses of the virus via computational virology methods.^9–22^ The transmission modes of the microdroplets^23,24^ enveloping the viruses can be classified in two. Direct transmission takes place when viruses are ”caught” airborne mainly via nose and mouth, while by indirect transmission, they are spread by touching surfaces with functional viruses and moving them into the respiratory system. This second mechanism can be very efficient, as suggested for the Wild-Type (WT) variant, that they can remain in certain types of surfaces for very prolonged periods of time, up to a few weeks.^4^ Based on this, the WHO has presented further recommendations on how to clean surfaces and on the continuous disinfection of hands. Due to the speed of the mutations of the SARS-CoV-2, the priority has always been placed on providing insights into the RBD-ACE2 interaction for the different mutations. This is a key step for the rapid development of new vaccines or therapies for SARS-CoV-2 mutations.^7^ In this context, the investigation of the interaction of other relevant VoCs with material surfaces has been lagging behind. In fact, little is known about how the delta and omicron variants interact with surfaces.

A computational characterization of the hydrophobic and hydrophilic interactions of the VoCs with different surfaces would provide biophysical insight into current experimental efforts made for developing immobilizable (filtering ^25^) and also virucide surfaces.^26^

All-atom molecular dynamics (MD) simulations can provide highly-valuable insight in the function of biological systems with high-resolution,^16,17,27–29^ and, in particular, on the interaction with different surfaces.^30–35^ From a molecular simulations viewpoint, the comparison of VoCs behaviour with different surfaces is still in its infancy. Pioneering studies on relevant hydrophobic, hydrophilic, skin, and coinage surfaces have been performed for the Wild-Type (WT) spike,^31–33,36^ and recent research focused only on the RBD interactions to very specific nanomaterials that can be degraded by macrophages ^34^. High-Speed AFM (HS-AFM) experiments have demonstrated the enhanced structural flexibility of the RBDs adsorption onto Mica substrates.^37^ However, the binding domain proteins (bottom of the RBD) at the bottom are currently arduous to image with HS-AFM techniques. Here, a particular challenge is tracking several binding domain mutations, ^38^ like the omicron variant, which highlights the urge to further elucidate their adsorption mechanisms onto hydrophobic and hydrophilic surfaces by computational biophysics techniques. In this work, we compare three Receptor Binding Domains (RBDs), namely those from the WT, Delta, and Omicron variants, interacting with two inanimate surfaces with the same structure and opposite polarities. To contextualize our findings, we contrast the interactions on inanimate surfaces with those on native-biological interfaces, specifically the ACE2 receptor. Our systematic study introduces simplified and polarizable surfaces, modeled in the shape of a molecular bilayer (Polarizable BL-PBL), and characterized by a contact angle that resembles hydrophobic/hydrophilic properties.^39–42^ In particular, we employ this bilayer model for completely hydrophobic and hydrophilic surfaces to characterize the interaction of different VoCs with substrates, and compare those results with the biological interaction of the cell surface receptor case (RBDs-ACE2). Including Glycans on top of the RBD proteins allowed us to determine their influence in the adsorption to inanimate bilayers, and explore their differences based on the VoCs. We chose the homogeneous polarizability (less specificity) scheme because it is a stepping stone towards a better understanding of the relationship between the RBD protein hydrophobicity/hydrophilicity mechanisms and their potential binding surfaces. Raising particular interest for surfaces with experimentally measured contact angles, which through nanoscale techniques, e.g. Scanning Probe Microscopy (SPM),^43,44^ broadens the application of this research to the co-design of functional materials and complements the electrostatic characterization of the RBDs.^15,21^ The evaluation of adsorption from our simulations covers the structural deformation, contact areas, contact histograms, single and group-based distance analysis, hydrogen bonding (hydrophilic surface), flexibility in 2D (Figure S15), and the formation of possible hydrophobic pockets for the interface sequences of 3 RBDs. The adsorption process of each RBD onto the hydrophobic and hydrophilic surfaces let us group the residues in two regions with the same amount of residues (see Figure 1, RBD legs in green (group 1) and orange (group 2), and Tables S7 and S8).

**Figure 1:**
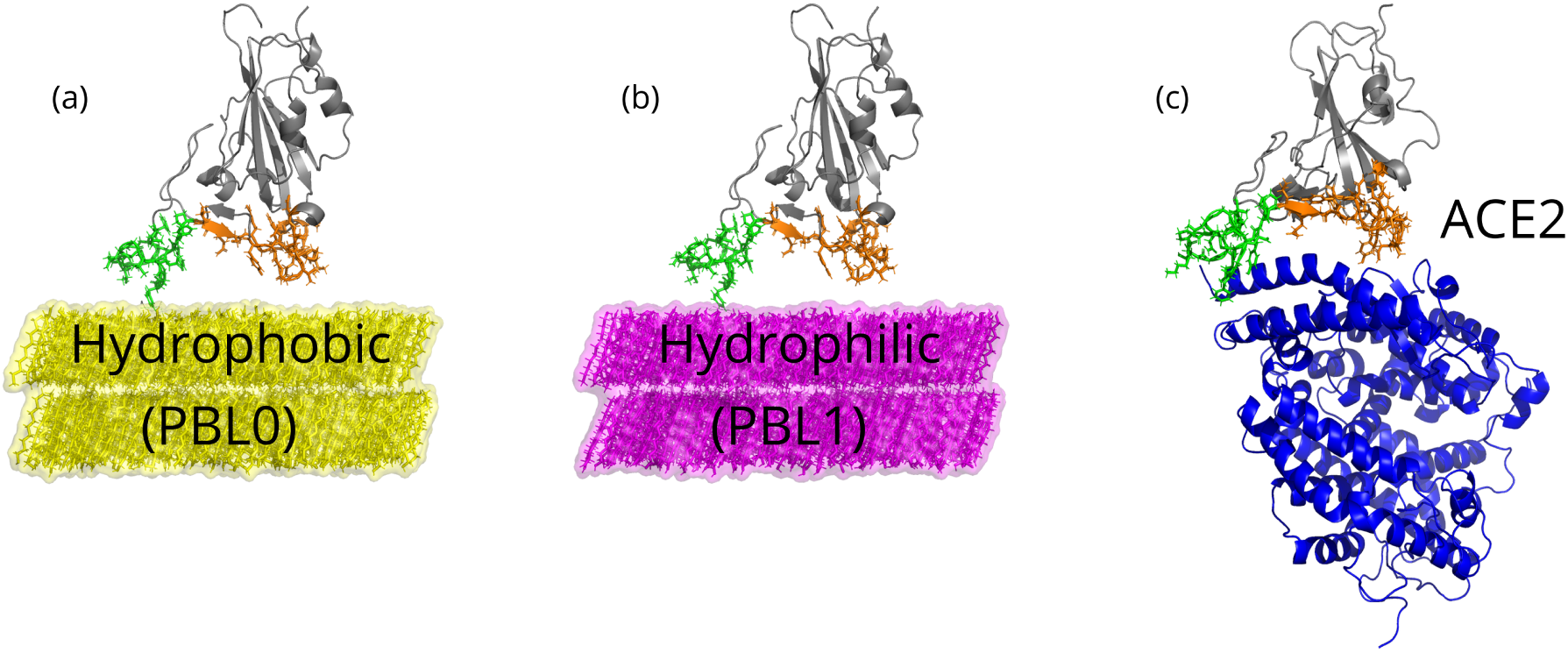
Snapshots of the three main scenarios simulated in this work. (a) Hydrophobic (PBLO) surface with an RBD, (b) hydrophilic(PBL1) surface with an RBD, and (c) the RBD-ACE2 complex, as a reference control model, are depicted. Note that in all cases, the left and right legs of the RBD are colored green and orange, respectively. Water molecules are removed for visualization reasons.

We found a fundamental difference in the adsorption of the RBDs onto inanimate (hydrophobic/hydrophilic) and biological surfaces, which highlights a flattening out mechanism of the RBDs over the bilayers, whereby the ACE2 interface remains far from reaching the polymer deformation observed at the bilayer interface. The comparison of contacts between the RBD-PBL0(hydrophobic) and RBD-PBL1(hydrophilic) is another analysis tool we combined with the contact area to show enhanced contact-areas and contacts of the PBL0 over PBL1. Strikingly, the RBDs including Glycans remain in similar limits as the purely proteinaceous RBDs, showing a minor contribution to adsorption of the polysaccharides in such interfaces. Again comparing the RBD residues that promote binding in the biological case with the inanimate surfaces, demonstrates that the adsorption mechanisms are controlled by residues in opposite regions of the RBM. The average distances of residues at the RBD regions for WT and delta show an unbalance landing of the RBM legs. On the contrary, the omicron variant reaches enhanced landing balance.

In general, more residues are closer to the tested surfaces for the WT variant, although the contact-area for the omicron variant is slightly wider than the WT one for the hydrophobic surface. This is related to the higher 2D-flexibility of the omicron adsorbing residues. While, for all variants, more residues are in contact with the hydrophobic surface than with the hydrophilic one, as previously reported for the WT variant.^31^ Furthermore, the mutations in omicron are precursors of possible hydrophobic-pocket like structures, which could stabilize its adsorption on the PBL0 surface.

## Results

### Morphological changes during the adsorption of the RBD onto polarized surfaces

Figures 2 and S1 show a collection of snapshots of the initial, mid and final adsorption stages of the RBD interacting with an hydrophobic (yellow) and hydrophilic (fuchsia in Figure S1) surfaces, respectively. Three views in each adsorption scenario capture the initial adsorption (1 ns) to the hydrophobic surface, namely, the proteinaceous RBD without Glycans (Figure 2 a,d,g), RBD with Glycans (Figure 2 b,e,h) and a rotated-RBD with Glycans (Figure 2 c,f,i) for the three VoCs. A bottom view of the same adsorption stage of the RBD protein is also shown in Figure S2. For the first and second columns, we clearly observe that both RBD legs (green and orange) are adsorbing onto the surface with slight preference on the green (Group 1). Interestingly, the mid column (with Glycans in vertical) show a Glycan chain (see also the Glycan structure in Figure S14) in red that is not directly interacting with the surface as the simulation starts. Hence, in order to further explore the Glycan contribution to the adsorption phenomenology of the RBDs, we rotated the molecule onto a extreme configuration, where the Glycan could form more contacts to the modeled surface. The latter shows that Glycans are also interacting to the hydrophobic surface, starting from the simulation genesis, however, due to its high flexibility^45^ is not remaining in contact to the surface.

**Figure 2:**
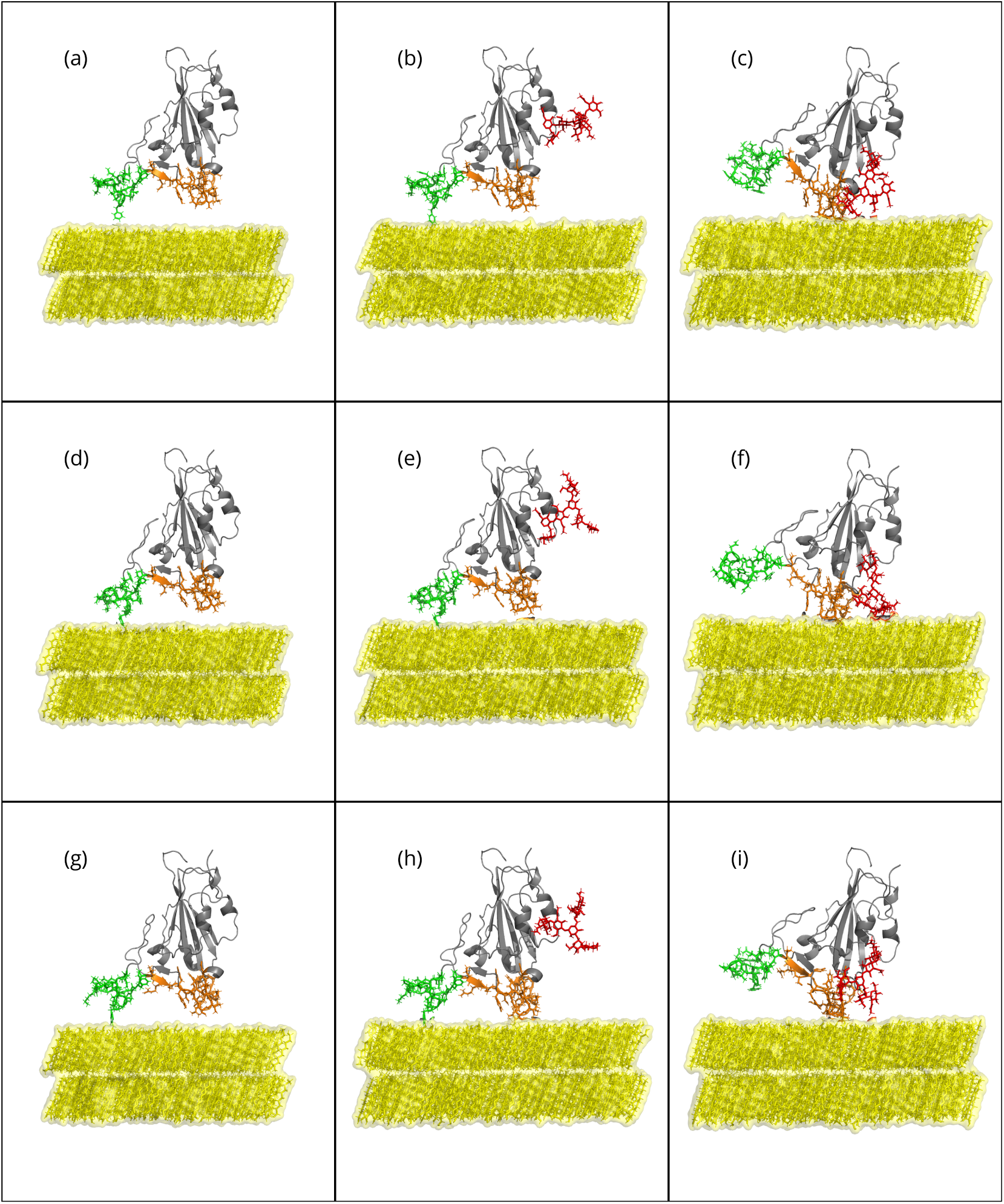
Side view snapshots of the RBD-PBL simulations performed for this research with the hydrophobic (PBL0) substrate at the beginning of the MD production. Rows show snapshots of the RBDs of (a-c) WT,(d-f) Delta, and (g-i) Omicron with the substrate alone, with its glycan standing vertically to the substrate, and rotated respect to the substrate with its glycan, from top to bottom.

The quantification of morphological changes in the RBM during adsorption are crucial steps to determining the difference in function of the protein due to structural deformation. Figure 3 shows the relationship between the radius of gyration *R*_g_*_⊥_* and *R*_g_*_∥_* of the RBM-inanimate (for both groups, as defined in Tables S7 and S8) and RBM-biological (both groups and the ACE2), and allows us a direct comparison among them for each VoC.^46–49^ The interfaces with the PBLs are roughly within the semiflexible polymer regime ([*R_g⊥_/R_g∥_*]^2^ *<*0.33).^50,51^ In contrast, the biological interface RBM-ACE2 shows a globular behavior. Al-though both hydrophobic and hydrophilic surfaces are in the same flexibility regime, we observe a difference in *R_g∥_* between Group 1 and Group 2, due to the initial conformation of the residues in Group 2, which are slightly more elongated (see Figures 3 (c) and (f) for initial structures between both groups). We highlight that the ratio of the [*R_g⊥_/R_g∥_*]^2^ reaches circa 3 times by comparing biological to biological-inanimate interfaces, which could trigger to irreversible deformations in the RBM as hypothesized elsewhere.^34^

**Figure 3:**
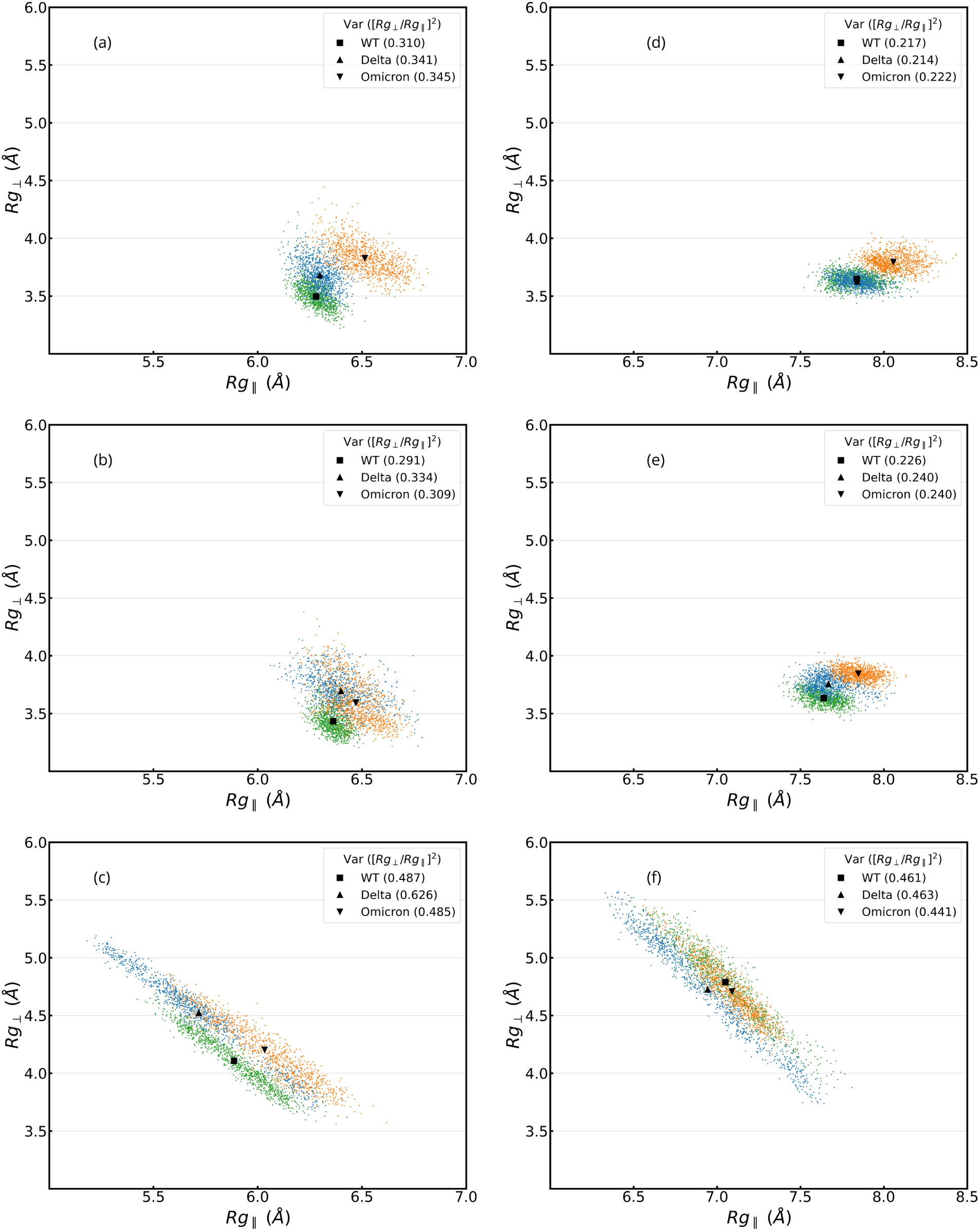
Perpendicular, *R_g⊥_*, vs. parallel radius gyration, *R_g∥_*, of the (a-c) group 1, and (d-f) group 2 of the RBD in presence of PBL0, PBL1 and ACE2 (from top to bottom). In each plot, green, blue, and orange dots are values for WT, Delta, and Omicron each 200ps, excluding the first 100ns. Black square, triangle and inverse triangle are the means of perpendicular and parallel radius gyrations, i.e. ⟨*R_g⊥_*⟩, and ⟨*R_g∥_*⟩. In the legends, [*R_g⊥_/R_g∥_*]^2^ ratio is shown in parenthesis for each case.

### Interaction between the RBDs and the polarized surfaces

Two additional physical quantities characterize the adsorption phenomena of the RBDs onto two antipodal surfaces. First, Figures 4(a) and (b), show the relationship between the number of contacts and the contact area for both surfaces, with and without Glycans. In particular, for the RBD-PBL0 interface has a gain in both total number of contacts and the contact areas in respect to the RBD-PBL1 interface (Figure 4(b)). Remarkably, the presence of Glycans has minor differences in both total contacts and contact areas, which lies within the standard deviations (see also Table S1). For the ACE2, the structures from crystallography provide an initial contact area that is larger than the other two surfaces(see Table S1). Note also, that the modeled polarizable surfaces do not contain Glycans, those are located only at the RBD. Another observation is the gain in contact area from the hydrophobic surface. This result goes inline with former observations comparing the interaction between the WT spike protein and the graphite and cellulose modeled surfaces,^31^ also shown in the distance analysis in the Distances and contact analysis section and the contact histograms of Figure 5. Most contact areas for the different VoC are in a similar range, with the exception of the omicron-PBL0 and omicron-Glycan-PBL1 interface with a notorious gain of around 1 *nm*^2^ in each case (Figure 4). Note also that the simulations on the PBL-1 are also prone to higher standard deviation due to hopping behaviour of the protein on hydrophilic surfaces (Figure 10(e) and (f)).

**Figure 4:**
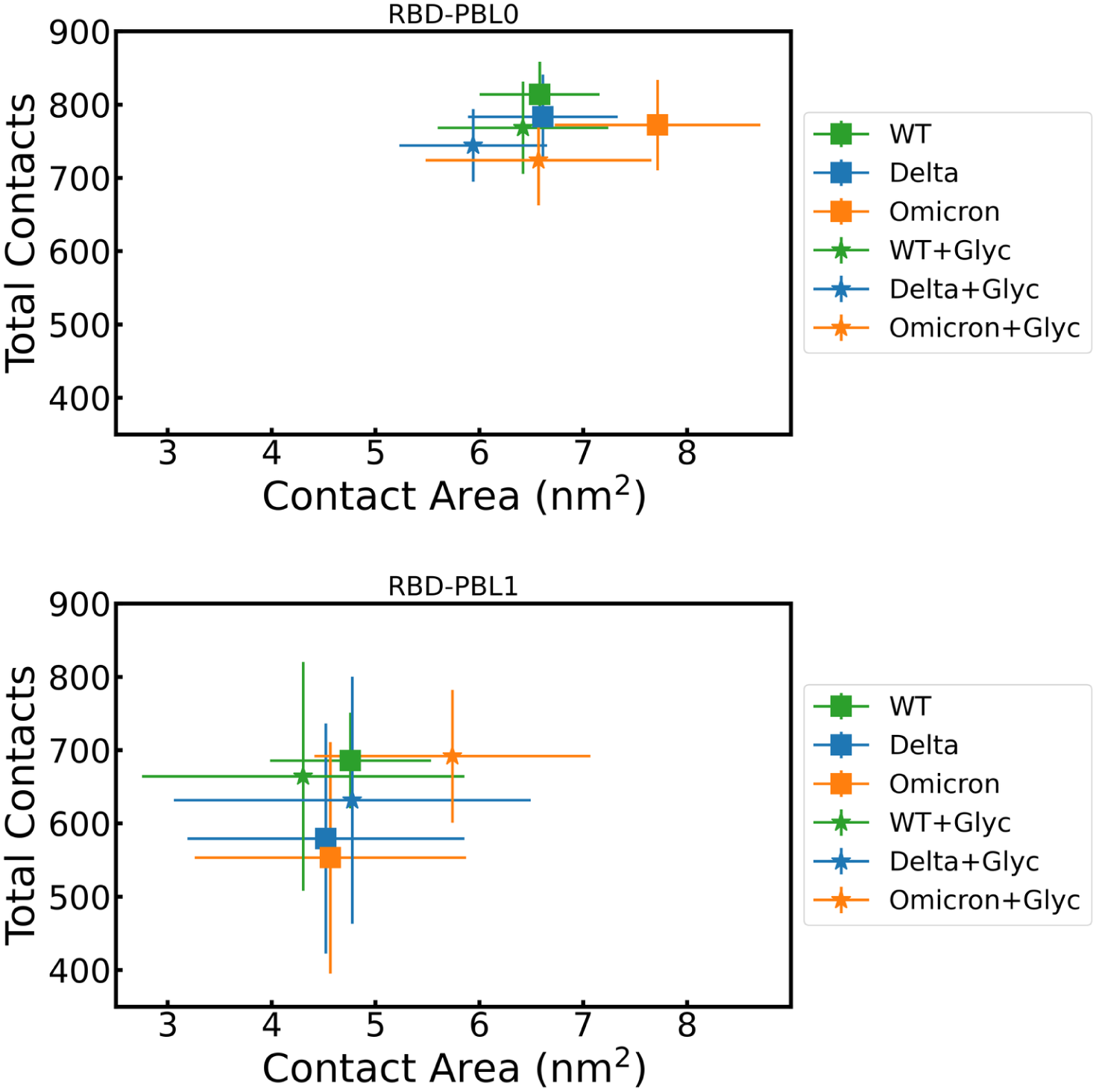
Total contacts per frame vs. contact area between the RBDs of WT (green), delta (blue), and omicron (orange) and the (top) hydrophobic (PBL0), and (bottom) hydrophilic (PBL1) surfaces. Squares are the mean over the whole trajectory of RBD-PBLs, and stars are the mean values over the whole trajectory of the simulations with glycan. Error bars show the standard deviation over time.

**Figure 5:**
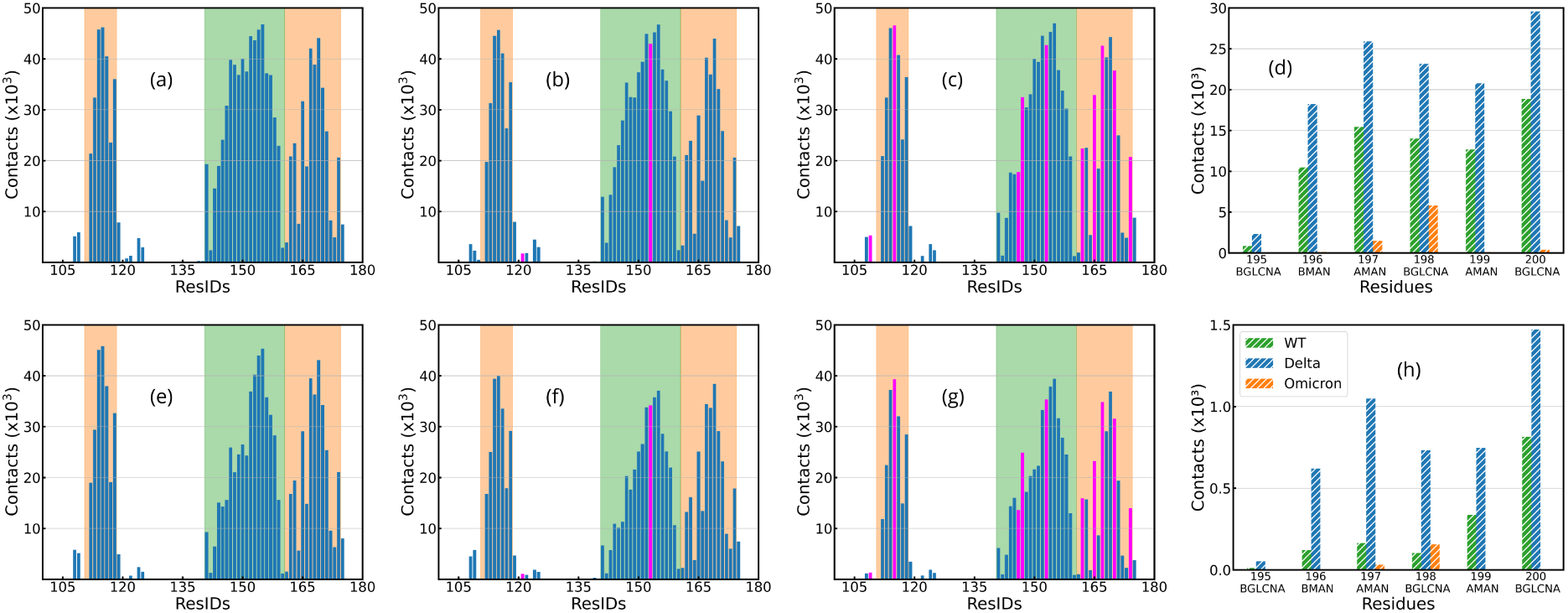
Residue contact histograms between the RBDs and the (a-d) PBL0, and the (e-h) PBL1 for (a,e) WT, (b,f) delta, and (c,g) omicron variants. Note that the mutated residues referenced to the WT are colored in fuchsia, and the legs of the RBD are shown by color in each region, green (left leg) and orange (right leg). (d) and (h) shows the contacts of each carbohydrate in the glycan to the hydrophobic and hydrophilic surfaces, respectively, in the rotated RBD-PBLs with glycans simulations, with the exception of 193-BGLCNA and 194-AFUC that have no contacts.

A second quantity is the number of contacts per residue, Figure 5(a-c) and (e-g) shows the accumulated number of contacts vs. RBD residues for the VoCs with both hydrophobic and hydrophilic surfaces. The mutated residues are highlighted in fuchsia and the RBD regions are also shadowed in green (left region-group 1) and orange (right region-group 2), as previously illustrated in Figure 1. The number of contacts in the hydrophobic interaction (Figures 5(a-c)) are more frequent than the hydrophilic surface(Figures 6(e-g)), as reported in the Distances and contact analysis Section. The RBD-ACE2 contacts can be found in Figure S3. In addition, Figure 5 (d) and (h) shows the accumulated number of contacts during the simulations with glycans (in the RBD rotated conformation) to the hydrophobic and hydrophilic surfaces, respectively. We observe a drastic decrease in terms of number of contacts compared to the protein side of the molecule, specially for the hydrophilic surface. As expected, we observed that most contacts between the glycans and the surface are at the tip of the glycan. However this interaction may be not only conducted by the enhanced flexibility of the Glycans, but also can be influenced by the mutations at the RBD. Complementary to the latter observation, we have analyzed local charges in the glycans, which may interact with the mutations in Group 2 of the omicron variant which contains more positive charges^29^ (see Figure S14).

**Figure 6:**
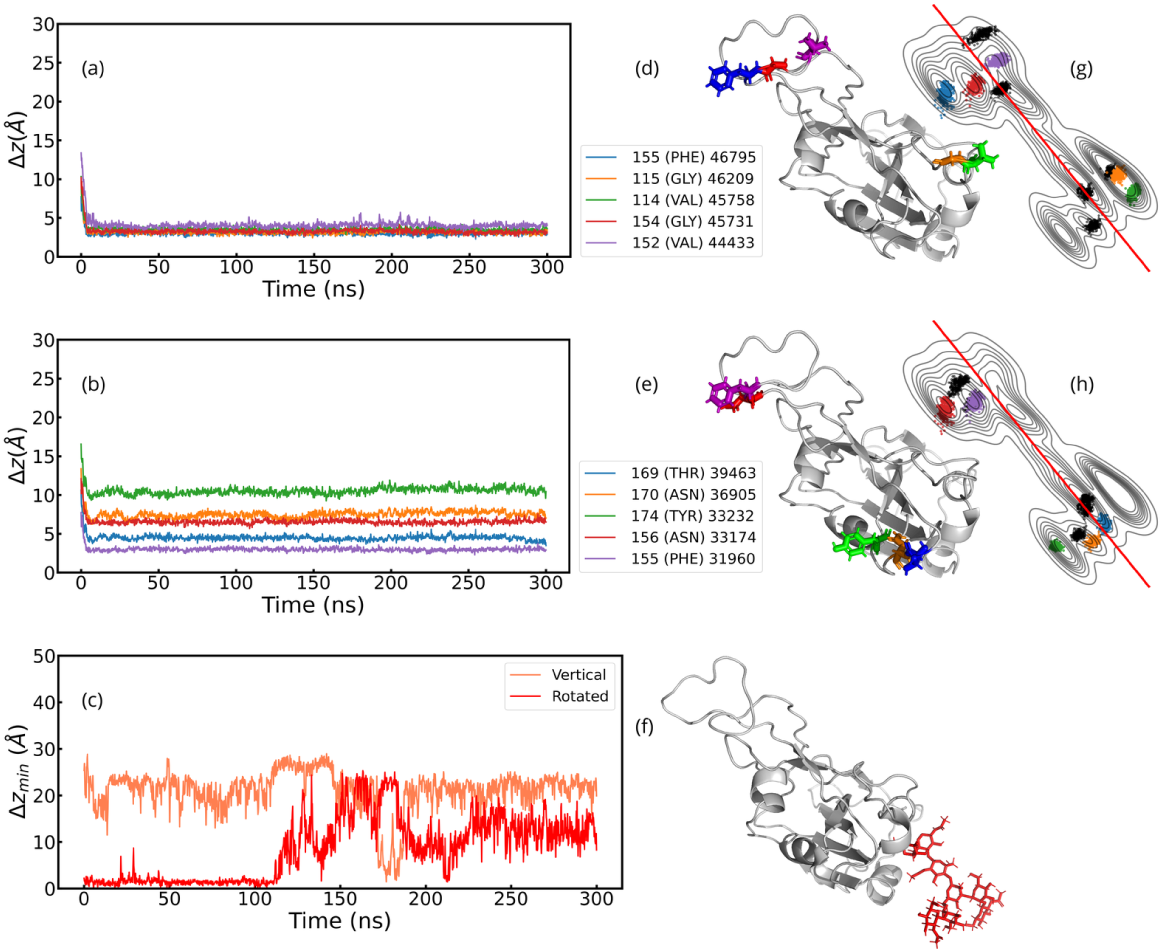
(a,b) Center of mass distance of the residues of WT-RBD to the hydrophobic substrate of the top 5 residues with most contacts with (a) the hydrophobic substrate and (b) the ACE2. Legends show the residue IDs, the residue names and the total contacts over the trajectory of each ranked residue (format: ResID (ResName) TotalContacts). Visualization of residues loci are also shown in (d) and (e) with colors corresponding with the distance plots (a-b). (c) shows the minimum distance between the glycan and the substrate in the hydrophobic surface in a vertical and rotated configuration. In (f), the glycan is shown in red. Note that all snapshots in this plot were taken from a bottom perspective. Complementary to (d) and (e), we present contour-line plots of top 10 residues with most contacts with (g) the hydrophobic substrate and (h) the ACE2. Note that in plots (g) and (h) the color code of the top 5 correspond to plots (d) and (e), the remaining 5 are depicted in black color.

### Distance and contact analysis of the RBDs and the polarized bilayers

Figures 6, 7, 8, 9, compile a detailed analysis of the WT and omicron RBDs, centered in the distances Δ*z* (as defined in the Methods Section) of the top 5 residues with most contacts in adsorption with the hydrophobic and hydrophilic surfaces (the case of the delta variant is also included in SI in Figures S4 and S5). These figures first display the distances curves and next the location of the residues in the RBD protein from a bottom perspective (right-hand side of those plots) The adsorption distance of the wild-type RBD onto the hydrophobic surface (Figures 6(a) and (b)) show only one coincident residue PHE-155, out of the total of 5 residues shown in the legend of Figure 6(a)). The ranks of the legends are given by the contact histograms of the PBL0 (Figure 5(a)) and the ACE2 (Figure 5(b)). The first presents substantial lower distances to the specific surface. These results are consistent with the contact histograms of Figure 5 and also aim to uncover the differences and coincidences between adsorption between the purely biological and biological-inanimate (biological-material) interface. In particular, performing a side-by-side analysis between the RBD-ACE2 and RBD-PBLs residues with more contacts. Strikingly, the residues loci of Figures 6(d) and (e), illustrate a notorious difference between the two groups of the RBM, where the adsorption (based on the top 5 analysis) in the case of the PBL0 is driven by the left region (Group 1), while the purely biological interface binding is rather triggered by Group 2. Note that the residues with more contacts (based on the histogram of Figure 5) highlight the higher proximity of the residues of the RBD-PBL0 interface with respect to the RBD-ACE2 interface. In addition, we provide KDE contour-line plots of top 10 residues with most contacts with (g) the hydrophobic substrate and the ACE2. Interestingly, by considering the closest 10 residues (Figures 6(a), (b) and Figures S6 (a),(b)) and comparing biological and biological-inanimate interfaces, a trend of contacts located in opposite regions at the Group 2 (referenced with the solid red line, see Figures 6(g) and (h)) is observed, which underlines the different landing-footprints on those two substrates. On top of this, it facilitates to understand the drastic morphological changes shown by exploring the different ratios [*R_g⊥_/R_g∥_*]^2^ onto both surfaces (see Figures 3(a), (d) and (c), (f)), by identifying a more spread residues *footprint* in the case of the flat PBL0. Furthermore, we also depicted the distances from the tip of the glycan to the PBL0 for both vertical and rotated configurations (see Figure 6(c)).

**Figure 7:**
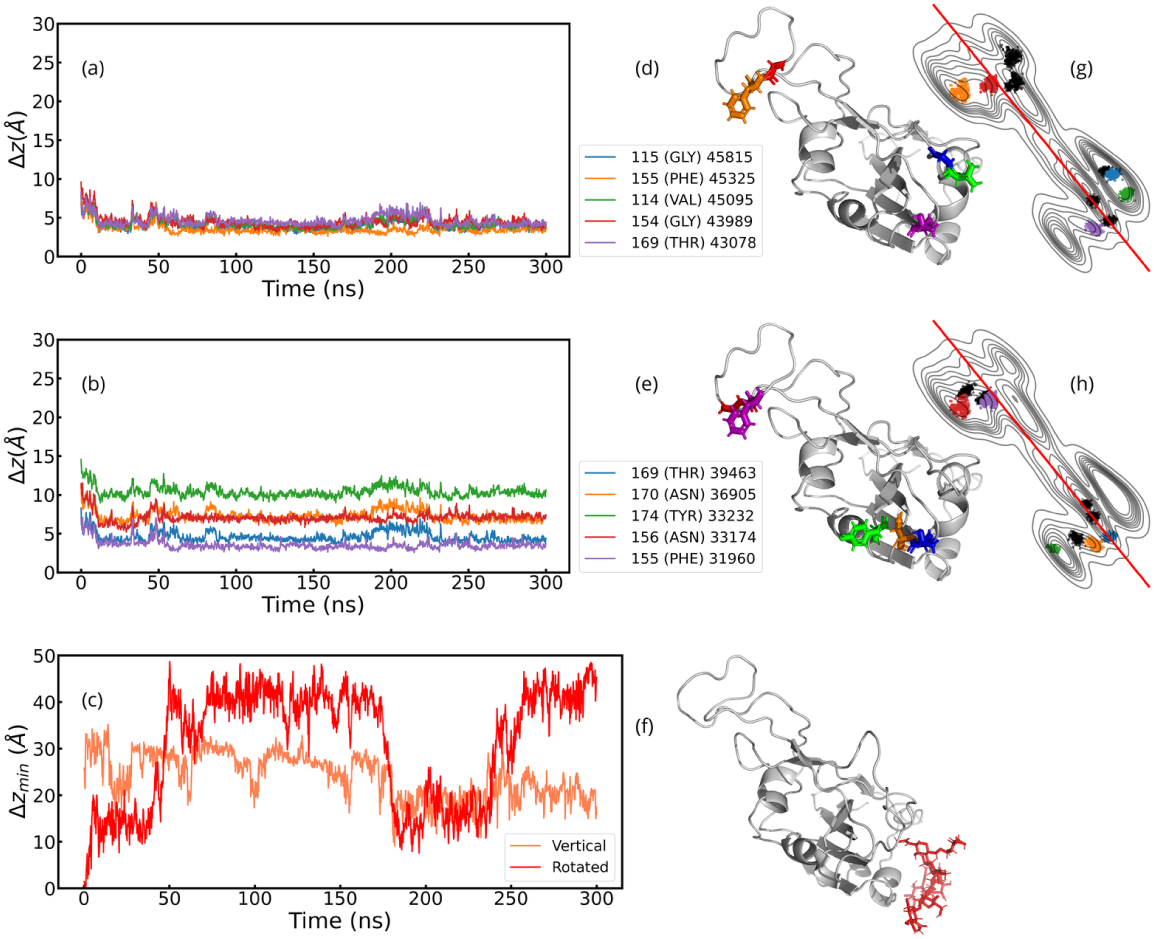
(a,b) Center of mass distance of the residues of WT-RBD to the hydrophilic substrate of the top 5 residues with most contacts with (a) the hydrophilic substrate and (b) the ACE2. Legends show the ResIDs, the residue names and the total contacts over the trajectory of each ranked residue (format: ResID (ResName) TotalContacts). Visualization of residues loci are also shown in (d) and (e) with colors corresponding with the distance plots (a-b). (c) shows the minimum distance between the glycan and the substrate in the hydrophilic surface in a vertical and rotated configuration. In (f), the glycan is shown in red. Note that all snapshots in this plot were taken from a bottom perspective. Complementary to (d) and (e), we present contour-line plots of top 10 residues with most contacts with (g) the hydrophobic substrate and (h) the ACE2. Note that in plots (g) and (h) the color code of the top 5 correspond to plots (d) and (e), the remaining 5 residues have adopted dark-grey color.

**Figure 8:**
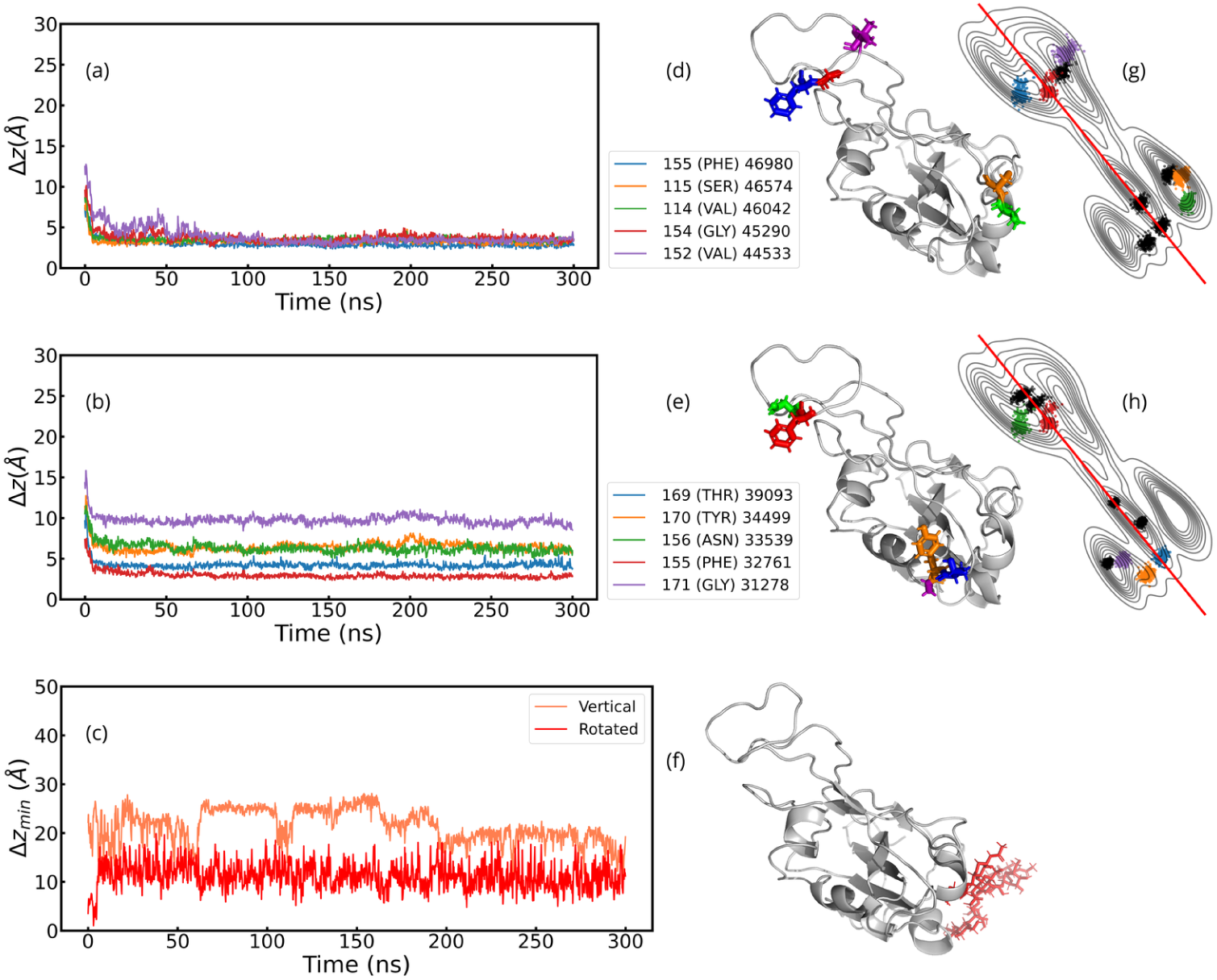
(a,b) Center of mass distance of the residues of Omicron-RBD to the hydrophobic substrate of the top 5 residues with most contacts with (a) the hydrophobic substrate and (b) the ACE2. Legends show the ResIDs, the residue names and the total contacts over the trajectory of each ranked residue (format: ResID (ResName) TotalContacts). Visualization of residues loci are also shown in (d) and (e) with colors corresponding with the distance plots (a-b). (c) shows the minimum distance between the glycan and the substrate in the hydrophobic surface in a vertical and rotated configuration. In (f), the glycan is shown in red. Note that all snapshots in this plot were taken from a bottom perspective. Complementary to (d) and (e), we present contour-line plots of top 10 residues with most contacts with (g) the hydrophobic substrate and (h) the ACE2. Note that in plots (g) and (h) the color code of the top 5 correspond to plots (d) and (e), the remaining 5 residues have adopted dark-grey color.

**Figure 9:**
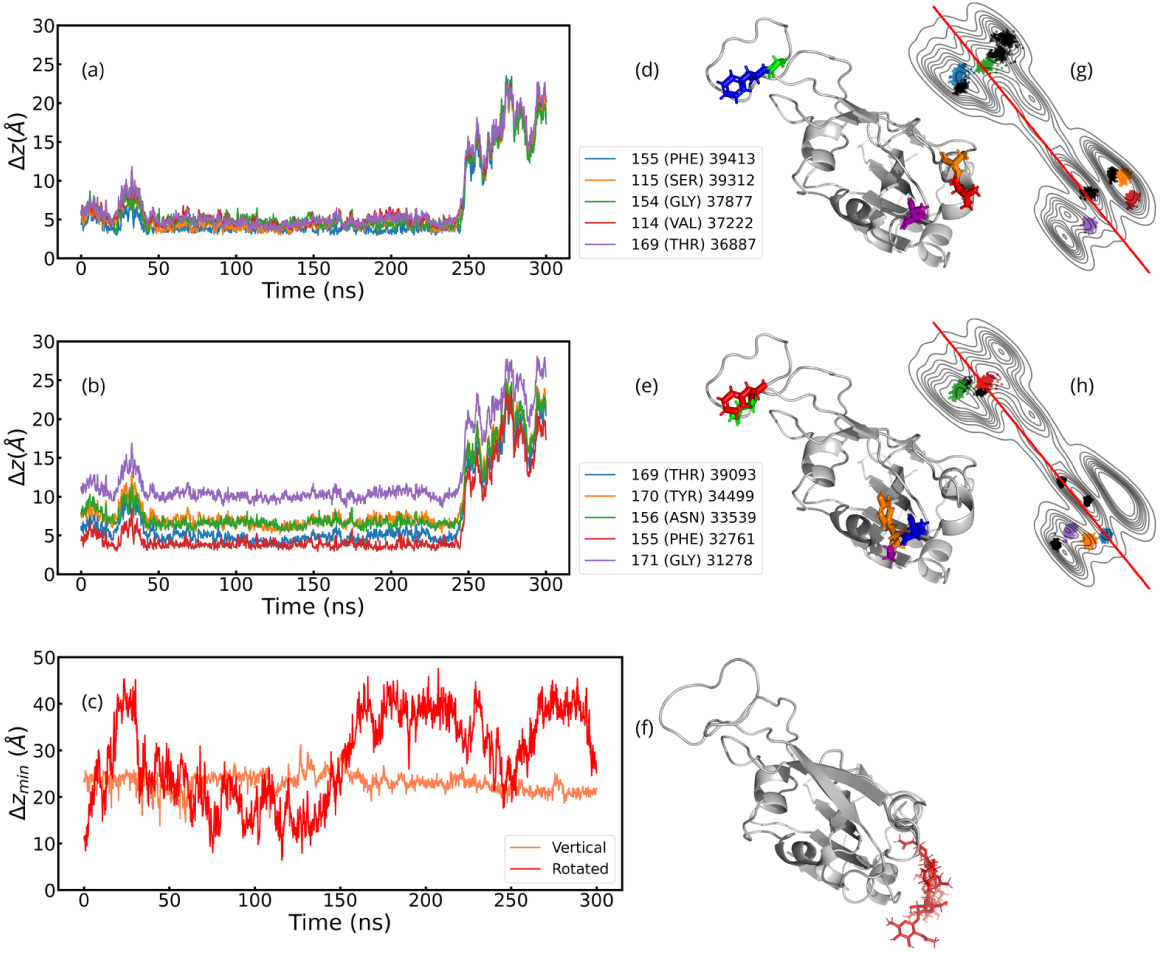
(a,b) Center of mass distance of the residues of Omicron-RBD to the hydrophilic substrate of the top 5 residues with most contacts with (a) the hydrophilic substrate and (b) the ACE2. Legends show the ResIDs, the residue names and the total contacts over the trajectory of each ranked residue (format: ResID (ResName) Total Contacts). Visualization of residues loci are also shown in (d) and (e) with colors corresponding with the distance plots (a-b). (c) shows the minimum distance between the glycan and the substrate in the hydrophilic surface in a vertical and rotated configuration. In (f), the glycan is shown in red. Note that all snapshots in this plot were taken from a bottom perspective. Complementary to (d) and (e), we present contour-line plots of top 10 residues with most contacts with (g) the hydrophobic substrate and (h) the ACE2. Note that in plots (g) and (h) the color code of the top 5 correspond to plots (d) and (e), the remaining 5 residues have adopted dark-grey color.

While for the hydrophilic surface the wild-type RBD adsorption tends to gain in fluctuations close to the surface (Figure 7). Such a slight oscillating behavior goes in line with the other variants (See Figures S5 and 9). For this surface the coincident residues during adsorption are again PHE-155 and THR-169, whereby based on the top-5 analysis, the right side of the RBD (Group 2) has 3 residues but two of them located in opposite corners within such regions. In general, the distances are higher than the hydrophobic case (Figure 6). This has been summarized in Table S2 of the SI. In addition, we provide contour-line plots of top 10 residues with most contacts with (g) the hydrophilic substrate and (h) the ACE2. Interestingly, by considering the closest 10 residues (Figures 7(a),(b) and Figures S7 (a),(b)) and comparing biological and biological-inanimate interfaces, a trend of opposite contacts for group 2 is observed, which underlines the different landing mechanisms. On top of this, it facilitate understanding the origin of the drastic morphological changes shown by exploring the different ratios [*R_g⊥_/R_g∥_*]^2^ on both interfaces (hydrophilic-PBL1 and ACE2) of Figures 3(b),(e) and (c),(f). Regarding glycans adsorption, the surface is not much attractive to the glycan, irrespective of its configuration (vertical or rotated).

The omicron variant shows a similar number of non-coincident contacts than the WT variant, in total 4 out of 5. However, the mutations here play a key role, for instance in the ACE-2 surface, by switching the tyrosine 174 (WT) by another one (ResIDs: 170 in omicron) but much closer to the other two top-5 residues, ResIDs: 169, 171 (see Figure 8(e)). This suggest a more specific and compact RBM interaction for the omicron variant, as also corroborated via extensive studies of the 3 variants binding energies.^15,21^ On the contrary, for the inanimate surface (see Figure 8(d)) the mutations within the top-5 residues in close contact do not show the binding-specificity. This alternative approach could bring potential new tools for modeling virucidal-surfaces and interpreting virus sensors, both are current applications requiring computational aided design of inanimate surfaces. In other words, designing surfaces by grouping different variants (non binding-specific), which could be based on a threshold of mutations per RBM groups or variant families.^52^ Certainly, our approach could be also used in the other direction for exploiting specificity, however, this will be part of a future work. By extending this benchmark to the top 10(See Figure S10), we found 2 mutated residues that start interacting with the PBL0 (see discussions in the Hydrogen bonding evolution Section). In addition, we provide contour-line plots of top 10 residues with most contacts with (g) the hydrophobic substrate and (h) the ACE2. By considering the closest 10 residues (Figures 8(a),(b) and Figures S10 (a),(b)) and comparing biological and biological-inanimate interfaces, a trend of opposite (referenced with the solid red line, see Figures 8(g) and (h)) contacts in group 2 is observed, which underlines the different landing-footprints. On top of this, it helps understanding the drastic morphological changes shown by exploring the ratio [*R_g⊥_/R_g∥_*]^2^ differences on both interfaces of Figures 3(a), (d) and (c), (f). The glycans at the omicron variant seem to fluctuate more in the 2D plane of the surface but maintaining a distance of ≈ 10 Å, which is not the case for WT (as depicted in the 2D-flexibility maps of figure S15).

Similar to the wild-type RBD, there is an oscillating (RBD-hopping) behavior in the adsorption process of the omicron RBD onto the hydrophilic surface (Figure 9). In this case, the oscillations are slightly larger that the ones observed for the WT variant. This was previously reported.^53^ The two coincident residues between RBD-PBL0 and RBD-ACE2 contacts are PHE-155 and THR-169, in both left and right regions respectively. This may indicate a balanced distribution of contacts while binding for both RBD legs. However, the final distribution of residues in contact shows differences between the remaining 3 residues in the top 5(Figure 9(d) and (e)). In addition, we provide contour-line plots of top 10 residues with most contacts in Figure 9(g) the hydrophilic substrate and Figure 9(h) the ACE2. Interestingly, by considering the closest 10 residues (Figures 9(a),(b) and Figures S11(a),(b)) and comparing biological and biological-inanimate interfaces, a trend of opposite contacts (referenced with the solid red line, see Figures 9(g) and (h)) is observed for group 2, which underlines the different landing-footprints. Group 1 has a particular loci of residues quite centralized at the left lobe (see Figures 9(h)). Similarly to previous cases (Figures 6, 7 and 8), it shows a much spread presence of contacts for the inanimate surface, in this case PBL1, which are related to the drastic differences in morphological changes shown by exploring the ratio [*R_g⊥_/R_g∥_*]^2^ on both interfaces (PBL1 and ACE2) of Figures 3(b),(e) and (c),(f). In the case of glycans we observe similar trends as for the WT case, where Glycans are not binding to the hydrophilic surface.

Complementary to the top 5 distance analysis, we tackled the distance distribution by both RBD regions defined in green (group 1) and orange(group 2) (Figure 1). For this, we use an average distance from all the residues, which at the same time is the distance of the residue’s center of mass, as defined in the Methods Section. Figure 10 shows the distances of the green (Group 1) and orange (Group 2)regions of the RBD for the WT, delta and omicron variants during their adsorption onto a hydrophobic and hydrophilic substrates. Singularly, both regions in the omicron RBD show similar and sometimes overlapping average distance between both regions (Figure 10(c) and its inset). This suggest a similar affinity onto the hydrophobic surface for both residues groups, which has been also shown as slightly decreased deformations for the *R*_g_*_⊥_* (see Figure 3). Another important aspect are the mutants, whereby Group 2 exhibits much more mutations than Group 1, which could explain the singular behavior for omicron in contrast with the WT and delta peers. For WT and delta on the hydrophobic substrate group 1 has a slight preference in terms of average distances. Such behavior is however, adaptable as shown for the omicron variant, where the adsorption process starts with slight preference for Group 2 and then after ≈ 80 ns Group 1 comes closer to the substrate, which could suggest an enhanced flexibility depending on the variant as recently reported.^28^ The hydrophilic substrates shows a different adsorption pattern, in both terms of increased average distances and also the hopping behaviour while landing (Figure 10(e) and (f)). Supporting information on the average values of the simulation trajectories shown in Figure 10 can be found in Table S2. Another complementary analysis to the top 5 residues includes the detailed next 5 residues, which builds the top 10 residues analysis in Figures S6-S11. This improves the understanding of the described hopping or balanced landing behavior during adsorption onto hydrophobic and hydrophilic substrates, beyond the contour-lines presented here.

**Figure 10:**
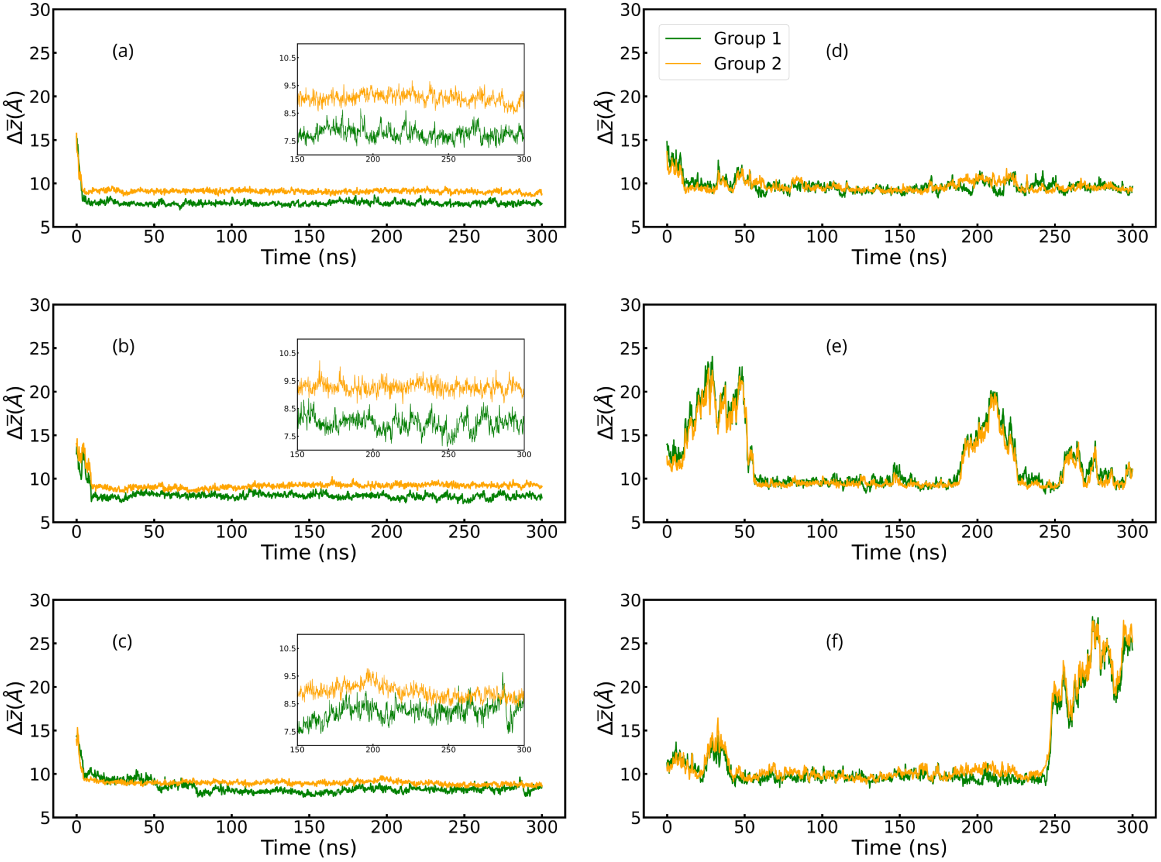
The average distance of the two regions of the RBDs, the left leg in green (Group 1) and the right leg in orange (Group 2) for (a-c) the hydrophobic, and (d-f) hydrophilic surfaces. Variants are ordered as Wild-Type, Delta, and Omicron, from top to bottom, i.e.(a) and (d) for WT, (b) and (e) for Delta, (c) and (f) for Omicron. Insets are a zoom of the trajectory from 150 to 300 ns.

Summarizing the distance analysis, we show in Figure 11 the residues adsorption at the interface are defined by distance thresholds of 6 Å (Figures 11(a) and (c)) and 10 Å (Figures 11(b) and (d)). As a general observation, we found more residues in the hydrophobic interfaces than in the hydrophilic ones. Moreover, the differences between variants for the hydrophobic surface are always around 7% (of the total number of residues), which shows a general similar adsorption process. However, this percentage changes once we look at the RBD legs (groups of residues), where omicron has 1 more residue placed closer to the surfaces than the WT or delta for group 2 (Figures 11(a)). And an opposite behaviour of omicron with less residues (1 less residue) at such a distance of this surface for group 1 compared to their peers (Figures 11(a)). At the 10 Å distance threshold (Figures 11(b)) omicron’s group 2 levels-off with delta and has only a minimal difference with WT. The landscape differs in group 1, where omicrons reaches ≈ 23% less residues than the WT and similarly with delta. Switching polarities to the hydrophilic substrate, we found a systematic (both for 6 Å and 10 Å distance thresholds) gain in residues for the WT variant over the other peers. In particular, in respect to the delta variant, reaching a 50% at the threshold 6 Å (half of the population form WT). While the second threshold 10 Å equalizes the difference between WT and delta. Now comparing numbers between hydrophobic and hydrophilic surfaces at the closest threshold of 6 Å, we can determine also ratio of 2 with more close contact residues at the hydrophobic interface. This might suggest a more reactive behavior onto hydrophilic surfaces due to the mutations at the RBM interface, which impedes the collective adsorption of the RBD to completely polar surfaces.A similar analysis has been previously performed. ^54^ An interconnected analysis to Figures 11(c) and (d) spots the Hydrogen bonds formed at the hydrophilic surface (see Hydrogen bonding evolution Section) and their distribution in the different RBDs footprint.

**Figure 11:**
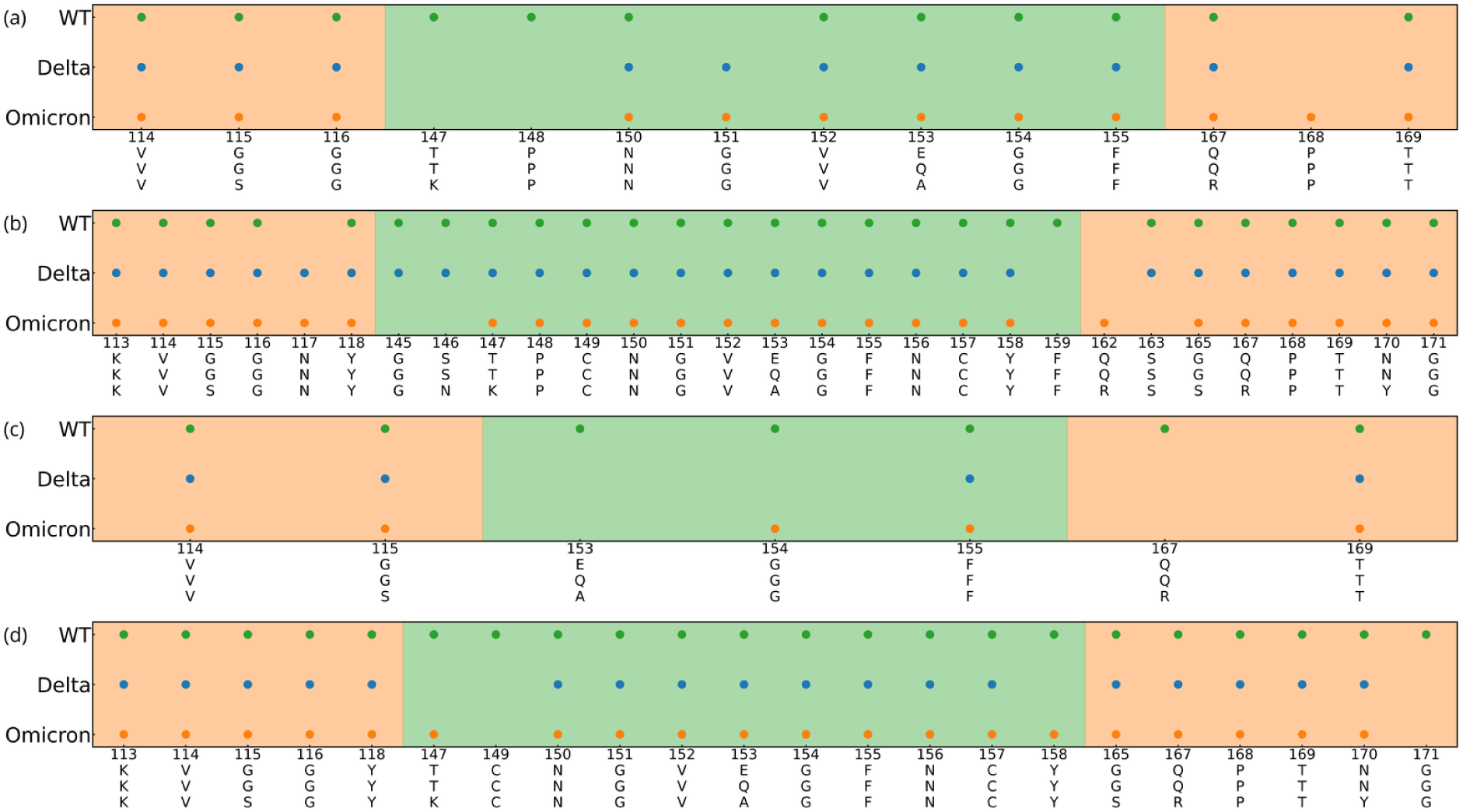
Residues with a average center of mass-surface distance smaller or equal to 6Å in (a) and (c), and 10Å in (b) and (d). (a-b) correspond to simulations with a hydrophobic surface and (c-d) to a hydrophilic surface. Note that all the distances plotted here are average distances of the total trajectory, excluding the first 100 ns, and distance values larger than 15Å.

To finalize this section, we analyze the ratios between the residues found at different interfaces to the surface (Table 1). Shows the ratio between the number of residues at the interface distances (6 Å, 10 Å and 14 Å) averaged out from the last 280ns of the trajectory (*N_avg_*) (see Figure 11), and *N_min_*the number of residues found at the interface at any time of the same trajectory, which fulfill the same distance criteria. The interfaces are defined by 3 threshold distances to the surface: 6Å, 10 Å and 14 Å. Table 1 shows the ratio *N_min_/N_avg_* at those 3 threshold distances, at the hydrophobic surface (left-side) and hydrophilic surface (right-side). In most of the cases, the following rule applies: the further the distance threshold the closer the values between *N_min_* and *N_avg_* and, hence,their ratio tend to the unity. Connecting the 3 threshold distances provides a sort of dynamic insight of the adsorption process, which can serve as input for the estimation of other distance-dependent type of interactions in max and average ranges, such as electrostatic forces ^18^ and van der Waals.^55–57^

**Table 1:**
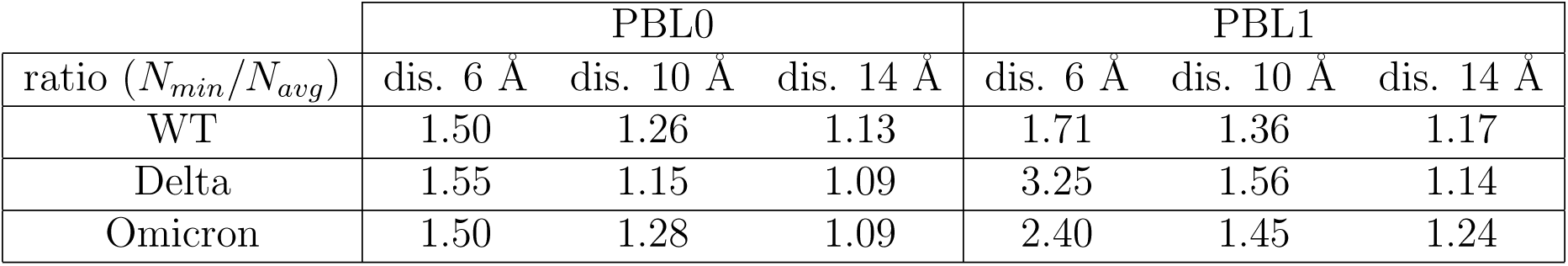
Ratios of the number of residues with minimum distance and the average distance, *N_min_/N_avg_*, for smaller or equal to average distances 6Å, 10Å and 14Å to the PBL0 (hydrophobic), and PBL1 (hydrophilic).

| ratio ($N_{min}/N_{avg}$) | PBL0 | | | PBL1 | | |
| --- | --- | --- | --- | --- | --- | --- |
|  | dis. 6 Å | dis. 10 Å | dis. 14 Å | dis. 6 Å | dis. 10 Å | dis. 14 Å |
| WT | 1.50 | 1.26 | 1.13 | 1.71 | 1.36 | 1.17 |
| Delta | 1.55 | 1.15 | 1.09 | 3.25 | 1.56 | 1.14 |
| Omicron | 1.50 | 1.28 | 1.09 | 2.40 | 1.45 | 1.24 |

At the same time, this analysis reinforces our previous description of a generalized intermittent contact of the RBD molecule onto hydrophilic surfaces, by observing that the overall sporadic closeness to the surface is much stronger for PBL1. Hence, the values in Table 1 are much higher for the hydrophilic surface. Remarkably, by using these ratios, we can also discern, at which distance the adsorption to the surface tends to reach a plateau. For example, which for PBL0 are clearly reached by the delta and omicron variants. At very close distances for PBL1 (6 Å, 10 Å and 14 Å), we observe that such a plateau is not reached, and the bouncing of residues is much higher at the closest distance of this analysis, as previously discussed in the distance analysis per RBDs regions (see Figure 11). This suggest that during adsorption at material surfaces there are more significant fluctuations than the other surfaces, and at the same time delta and omicron are pointed-out as more reactive ones.

Connected to Table 1, we present Tables S5 and S6, which show the *N_min_/N_avg_* normalized by the maximum number of contacts and Tables S3 and S4 show those normalization factors.

### Hydrogen bonding evolution and hydrophobic formations at the interfaces RBD with polarized surfaces

In this section we show first the Hydrogen bonds formed at the interface between the three VoCs and the hydrophilic surface as averages over time. Figure 12, shows for all residues that form H-bonds per variant of concern. Here, we rapidly identify the location of the H-bonds by including the contour-lines of the different RBDs. Strikingly, the mutations of residues form hydrophobic to hydrophilic in the omicron variant are rapidly spotted in group 2 (right side of the plot), reaching ≈ 70% of the total H-bonds. Also, the mutation in ResID 153 from WT (GLU) to delta (GLN) decreases the presence of H-bonding, which are consistent with the literature.^58^ In terms of H-bonds, the WT variant is almost perfectly balanced by 50% in both RBM regions (left and right). The ResIDs found in the top-10 distance analysis are also found in the H-bonds criteria. Similar analysis for the WT variant have been also reported the presence of H-Bonds with hydrophilic surfaces. ^31^

**Figure 12:**
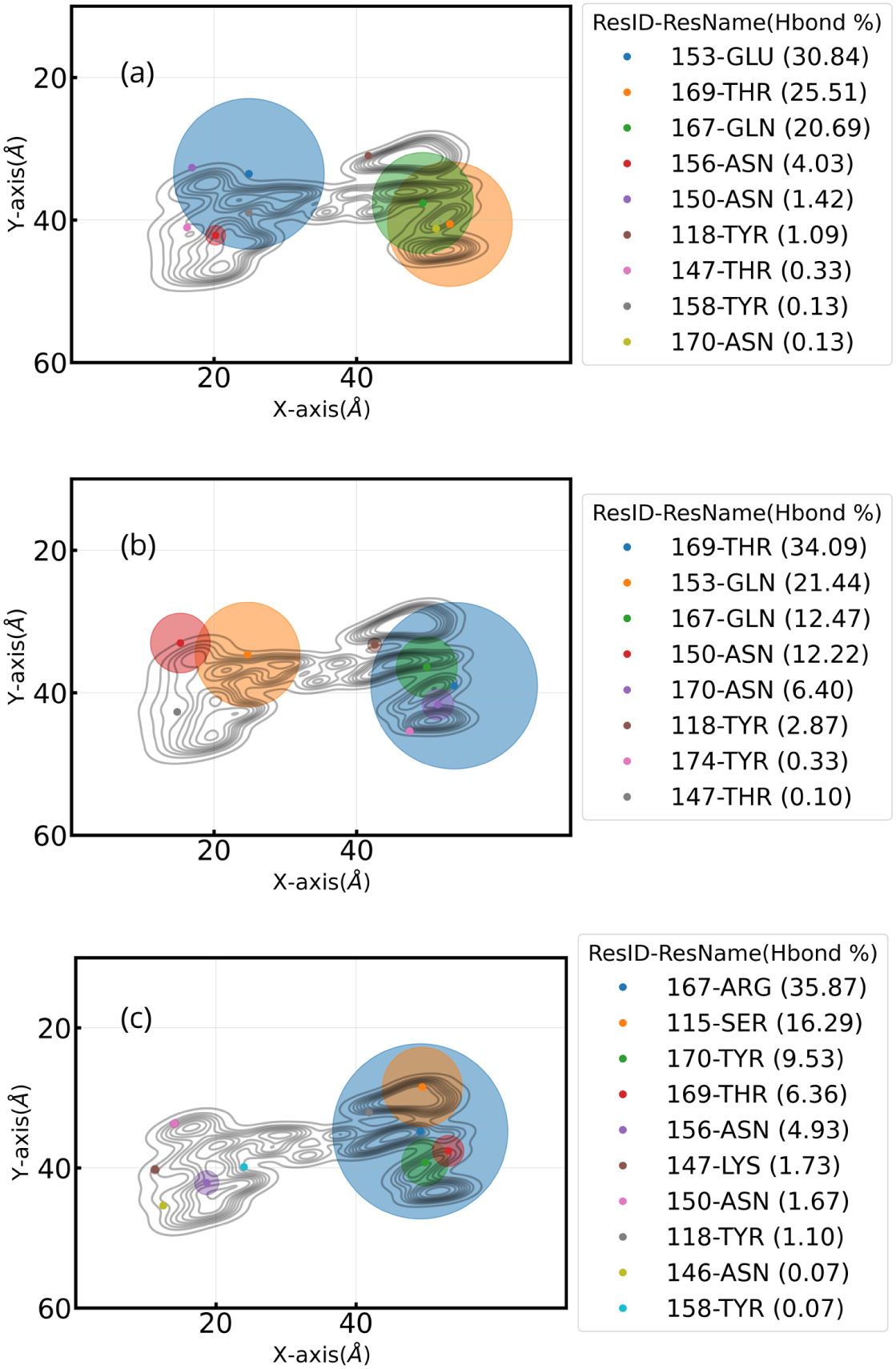
H-bonds of residues with hydrophilic surface for (a) WT, (b) Delta, and (c) Omicron. Scatter plot shows location of all center of mass of hydrophilic residues that have H-bonds with the surface from a top view. Circles are representations the percentage of the trajectory in which each residue have H-bonds. Scatter color vary according to the ranking of this percentage. In the legend, in parenthesis next to the ResIDs and residue names, the percentage of time with H-bonds is shown for each residues is shown. Note that all plots here include the contour-line plots.

After noticing that the RBM’s group 1 is the most flexible region. We analyzed the formation of ring-like structures and in perspective to the distance from the surface, rather crook-handle like structures. In Figure 13(d), we show those formations for the different variants and the hydrophobic surface represented by a collective variable *d_pocket_*((see Figure 13(e))) as the simulation time evolves. Remarkably, the residue 153 mutates from hydrophilic (in WT-GLU and delta-GLN) to hydrophobic (in omicron ALA) and allows the formation of hydrophobic pockets which closes the crook-handle formation (see Figure 13(c)). This hydrophobic formations could also drive the further design of rather nano-patterned hydrophobic surfaces to trap in principle only one specific variant, as discussed in the Distances and contact analysis Section.

**Figure 13:**
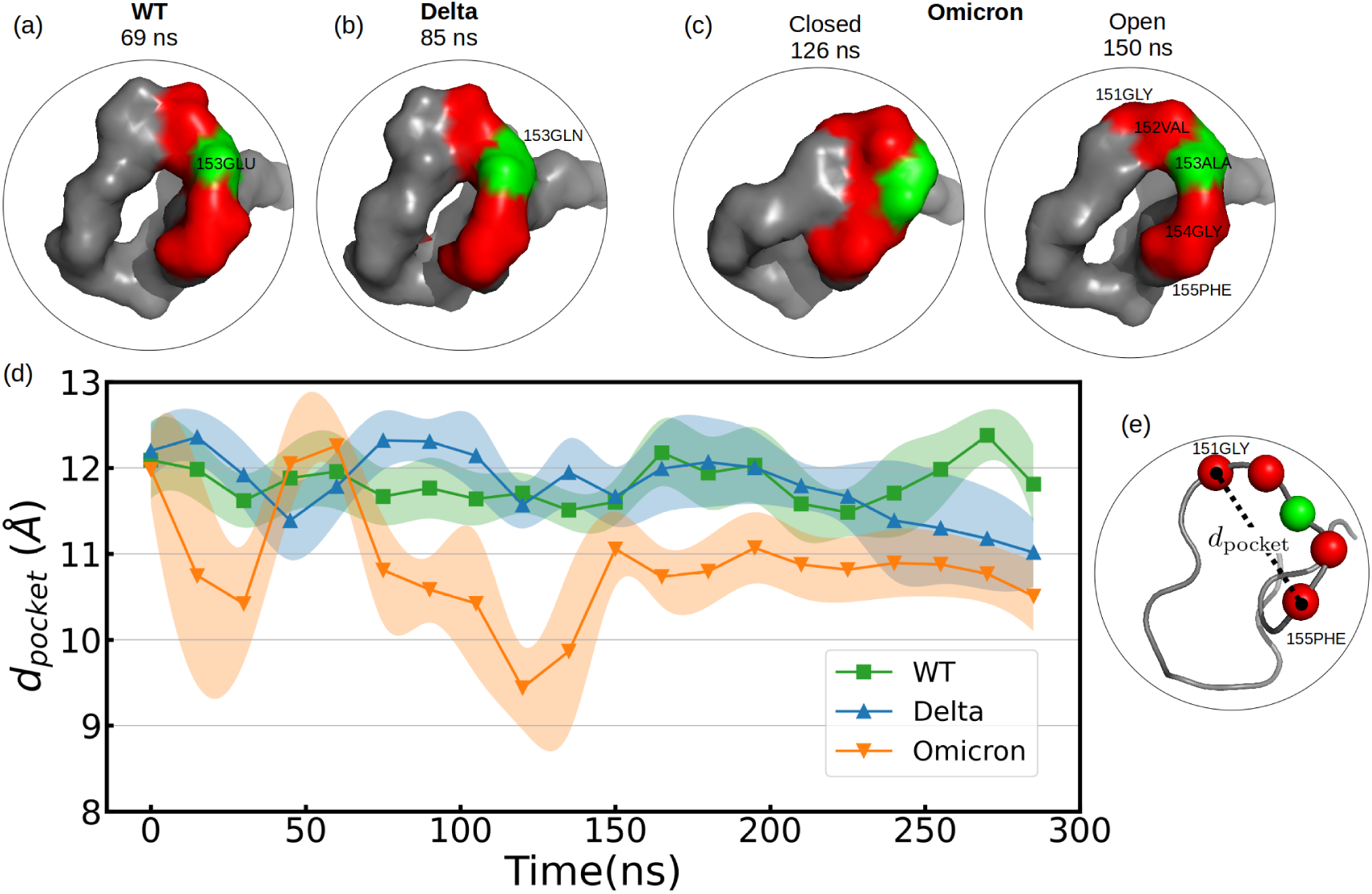
(a-c) Crook-handle formation in group 1 in presence of an hydrophobic surface. In red, the loci of the C-alpha atoms corresponding to the residues 151, 152, 154 and 155 is represented; in green is residue 153, which mutates from WT to the other two VoCs. Residue 153 is part of the crook-handle, forming a hydrophobic pocket in Omicron. (d) The mean C-alpha atoms distance between residues 151-GLY and 155-PHE over each 15ns, *d_pocket_*, which is depicted in (e). Colored shadows represents the standard error over these 15 ns. Note that the perimeter of the crook-handle from residue 151-GLY to 155-PHE was 15.3 ± 0.1Å for all variants. In omicron a breathing mechanism is promoted (see in (c) that at 126ns closed state, and open at 150ns) due to attraction of 153-ALA to the hydrophobic surface. The adsorption of 153-ALA to the PBL0, forces 154-GLY to be reoriented to the 149-CYS direction for geometrical reasons.

## Discussion

We have presented extensive molecular dynamics simulations of three SARS-CoV-2 receptor binding domains with two different model surfaces. These simplified surfaces have been implemented in order to distinguish the adsorption processes of the WT, delta and omicron RBDs driven by homogeneous hydrophobic and hydrophilic surfaces. In addition, as a reference to determine the ’biological contacts’ of the RBD we simulated RBD-ACE2 interfaces of the tackled variants. Initially, the RBDs adsorb to both surfaces through residues, which are mostly located in the RBM region. From this point, the adsorption on each surface describes substantially different patterns. Overall, the RBDs adsorption onto hydrophobic surfaces show a gain in number of residues at the interface, contact area and contact histograms, compared to the hydrophilic one. However, in detail each RBD variant presents complex and different surface-adsorption mechanisms. Considering two regions of adsorption at the RBM-surface interface with the same total amount of residues, we show how the regions balances out (hops) during adsorption and favors the contact formation. Specifically, the WT-RBD and hydrophobic-surface presents an enhanced adsorption than the other two peers, quantified in terms of the average distances of the RBM-surface, and contact histograms. While for the hydrophilic surface, WT is also presenting better adsorption, as quantified by both the average distances of the RBM-surface and the location of the H-bonds distribute along the whole RBM contact-region. In contrast, the adsorption onto hydrophilic surfaces for delta and omicron show intermittent contact, where the H-bonds are mostly distributed in one end of the RBM, namely, group two (right-leg). The average group distances shows for both delta and omicron a distinguishable oscillatory behaviour onto hydrophilic surfaces, specially for omicron where the H-bonds are 90% localized in the RBM’s group 2 resulting in a hinged-oscillatory mechanism. Consistent with recent structural biology insights in the literature.^59^

The simulations were also used to compare adsorption trends of the residues in contact between the RBDs-ACE2 (biological contacts) and RBDs-PBLs(inanimate/material interfaces). Such an analysis provides important differences between the preferred adsorption regions of the RBM, depending on the biological (specific) and the inanimate (non-specific purely hydrophobic or hydrophilic surfaces). Our results show systematically that the RBM’s group 2(right leg) adsorption takes place in residues of opposite domains depending on the biological or non-biological origin of the surfaces. In the RBM’s group 1, there are also changes in the adsorption footprints between inanimate and biological surfaces, however, they are not as systematic as in group 2. In fact, those differences in landing-footprints can also explain the RBDs enhanced adsorption-driven deformation, which exhibits deformation ratios of [*R_g⊥_/R_g∥_*]^2^ with differences in the order of ≈ twofold, where the flat hydrophobic and hydrophilic surfaces result in higher deformations of the protein.

We demonstrated different landing-footprints and deformation mechanisms of the mutations in the RBDs proteins that can be directly applied for the correct interpretation of experiments, similarly to the recent on high-speed AFM work performed for RBDs to ACE-2.^37^ Future assessments for the design of virucidal surfaces with novel high-speed AFM techniques are also warranted through collaborations. It is evident to note that our study has to be treated as a first step in the understanding of the molecular adsorption between VoCs and surfaces. Nonetheless, it sheds light onto the complex adsorption-driven deformation of the proteins, by considering hydrophobic and hydrophilic interactions, which are key to understand short-range binding. Strikingly, the RBM is capable of deforming up to two times more at a surface with only minor changes in their secondary structure (see Figure S13), suggest suggesting a shock-absorbing mechanism. Beyond the scope of this article, our results can be combined with current multiscale models of the whole virus ^19,20^ to tackle the shock-absorbing feature of the RBM. In a nutshell, a predetermined set of residues in group 2 facing adsorption only in one loci rapidly switch to the opposite region once the biological surface turns into one of the modeled inanimate ones. This adsorption assessment, highlights differences from biological to inanimate surfaces, such as their twofold deformation, which provides crucial complementary insights to the current experimental development of virucidal-surfaces and biosensors.

## Methods

### Molecular Dynamics

The simulations in this research do not consider the whole spike for the VoCs, because it has been shown in the WT that most interactions with surfaces were identified at the RBD,^31^ more specifically in the Receptor Binding Motif (RBM), which is the region of the spike protein interfaced to the ACE2.^13^ We explicitly include the RBD in the simulations and mimic the interaction of the RBD with the rest of the spike by introducing some mechanical constraints at the contacts between the RBD and the S1 region, as explained later in this Section.

All-atom simulations were carried out with Gromacs 2023^60^ and the system components (protein, ions, and the polarizable bilayer) were modeled using CHARMM36^61,62^ forcefield, and TIP3P^63^ for the water. CHARMM-GUI was used to join the glycan to the RBDs. Energy minimization used CPUs, while all production runs used 1xGPU, as the former scaled better than 2xGPUs for our systems. All RBD models for the initial configurations were adopted from previously published results by,^15^ where we added the disulfide bonds to the VoCs. The hydrophobic (PBL0) and hydrophilic (PBL1) surfaces were built from a small patch of decanol (DOL), in which restraints were used in order to maintain the bilayer shape and avoid any effects of the mechanical properties of the surface, as discussed in previous works,^39–41^ the bilayer model does not include any curvature(is not flexible) and also no molecular defects. The replicated PBL (using *gmx editconf* ) patch was solvated in a slab formed water box. The RBD models for WT, Delta, and Omicron were added to a 8nm×8nm×12nm cubic box containing the bilayers. The OH-groups of the DOL chains were tuned to 0 or 1 for hydrophobic and hydrophilic surfaces, respectively. ^41^ This procedure was repeated 3 times for each polarity, aggregating from 3 replicas per VoCs (WT, delta and omicron) obtaining 18 configurations. The same procedure was followed for the RBDs with ACE2 also using cubic boxes. All simulations included ions to work under neutral charge conditions. Periodic boundary conditions were applied and PME was used for long-range electrostatics. Minimization was done by steepest descent (50000 steps) with integration steps of 0.01 ps. The equilibration time for the NVT and NPT was 100ps, respectively. For each RBD, we determined the contacts between the RBD and the S1 region. Those contacts were applied as position restraints of 250 kJ mol^-1^ nm^-2^ in x and y axes. In other words, we considered as flexible regions of the RBD all the others that are not in contact to the S1. However, in order to quantify adsorption, we kept the z-axis free of restraints in all RBD-S1 contacts. Note also that the position restraints do not apply to the RBM interface (See also Figure S12 and Table S10). Production simulations began from the final equilibrated snapshots, and three copies (with angular rotations of up to 3 degrees from the reference) of each system were simulated. Finally, 300 ns of trajectories of each replica were collected as described in Table 2. Note that all production runs used an integration step of 2 fs. Starting configurations from the MD production can be found in a Zenodo repository.^64^

**Table 2:** The table contains the general configurations of the simulated systems. Note that 576 decanol molecules were used in systems without glycans, and 648 decanol molecules in simulations with glycans. Simulations of RBD-PBLs with glycans are done in two different configurations of the RBD respect to the PBL; a vertical RBD (1v) and a rotated RBD(1r) configuration. Snapshots of the initial configurations for PBL0 are shown in Fig.2.

| VoC | No. of replicas | PBL polarities | Box size (nm<sup>3</sup>) | Atoms | Time per replica [ns] |
| --- | --- | --- | --- | --- | --- |
| WT | 3 | 2 | 7.5×7.5×12 | 2980 | 300 |
| Delta | 3 | 2 | 7.5×7.5×12 | 2987 | 300 |
| Omicron | 3 | 2 | 7.5×7.5×12 | 3036 | 300 |
| WT+Glyc | 2 (1r+1v) | 2 | 8.5×7.6×12 | 2979+192 | 300 |
| Delta+Glyc | 2 (1r+1v) | 2 | 8.5×7.6×12 | 2986+192 | 300 |
| Omicron+Glyc | 2 (1r+1v) | 2 | 8.5×7.6×12 | 3035+192 | 300 |
| WT+ACE2 | 3 | - | 15.2×15.2×15.2 | 12510 | 300 |
| Delta+ACE2 | 3 | - | 15.2×15.2×15.2 | 12523 | 300 |
| Omicron+ACE2 | 3 | - | 15.2×15.2×15.2 | 12564 | 300 |

### Structural analysis

In all cases, we have calculated the Radius of Gyration parallel (*R*_g_*_∥_*) and perpendicular (*R*_g_*_⊥_*) were calculated using the equations as follows,

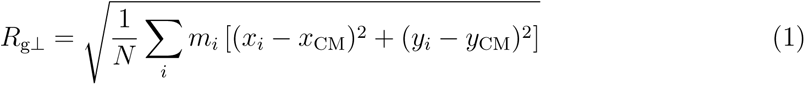

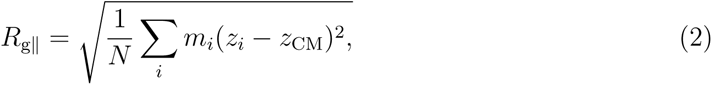

where **R**_CM_ = (*x*_CM_, *y*_CM_, *z*_CM_) is the position of the center of mass, *m_i_*the mass of each residue and *N* the total number of residues.

Moreover, the contact area has been calculated by subtracting (*SASA_RBD_* +*SASA_PBL_* −*SASA_RBD_*_+_*_PBL_*)*/*2 and (*SASA_RBD_* + *SASA_ACE_*_2_ − *SASA_RBD_*_+_*_ACE_*_2_)*/*2, for the PBL and ACE2 simulations, respectively.

### Contacts analysis

Two types of contact analysis have been considered, one using the No. of Contacts vs. simulation time and the other No. of Contacts vs. Residues. The latter includes the highest resolution trajectory available and sums up the contacts found with the PBL. While the No. of Contacts vs. simulation time includes temporal averages of the contacts every 1 ns, this facilitates the comparison with the distance analysis presented in the next paragraph. The cutoff we used in all cases is 14Å. To this matter, Contacts were counted at each frame when distances of the center of mass of residues to the surfaces (both PBLs or the ACE2) were less than 14Å.

### Distance analysis

Given the simplified definition of the polarizable bilayer, we provide an in-house distance analysis that shows the explicit distance of the center of mass to the PBL, according to,

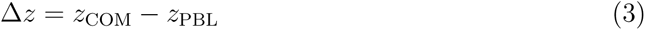

where *z*_COM_ is the position of the center of mass of each residue and *z*_PBL_ is the position of the bilayer. This quantity is calculated every 200 ps, once per snapshot.

## Author Contributions

H.V.G. conceived the research; H.V.G. and R. P. designed the research; A.B. performed the molecular dynamics simulations; H.V.G. and A.B. performed basic and advanced analysis; A.B., H.V.G. and R.P. interpreted the data; H.V.G and R.P. wrote the manuscript and supervised the research. All authors have read and agreed to the published version of the manuscript.

## Acknowledgement

We thank Matej Kanduč for illuminating discussions on the molecular modeling of a polarizable surface, Prof. R. Amaro for advice regarding the modeling of Glycans. The authors acknowledge the financial support of the Comunidad de Madrid and the European Union (VIRMAT REACT-UE project) through the European Regional Development Fund (ERDF), financed as part of the Union response to COVID-19 pandemic. H.V.G thanks the financial support by the Slovenian Research Agency (Funding No. P1-0055) and. R.P. acknowledges support from the Spanish Ministry of Science and Innovation, through project PID2020-115864RB-I00 and the “Maŕıa de Maeztu” Programme for Units of Excellence in R&D (CEX2018-000805-M). We thank the Red Española de Supercomputacíon (RES) for the computing time and technical support at the Finisterrae III supercomputer projects FI-2023-1-0029 and FI-2023-2-0036.

## Supplementary information

The Supporting Information available within this article contains: The RBD-PBL snapshots for the hydrophilic substrate; Bottom view snapshots of the RBDs adsorbed onto both substrates. Mean values with STD of the contact area calculations. The contact histograms of the RBD-ACE2 interactions. Top 5 COM distances for the delta variant onto both hydrophilic and hydrophobic surfaces; Top 5 COM distances for the 3 variants onto both hydrophilic and hydrophobic surfaces. Tables containing the residues involved in Groups 1 and 2 for the analysis; An illustration of the position restraints applied to all RBDs and the corresponding table. Furthermore, a plot containing the distributions of secondary structures for the RBD alone and RBDs onto modeled surfaces.

## Data and Software Availability

All-atom simulations were carried out with Gromacs 2023; the corresponding parameter files, input files, topologies, position restraints, and initial configurations, as well as, the analysis scripts as jupyter notebooks and data produced in this work are available on the Zenodo repository. https://zenodo.org/records/10479755.

