## Supplementary Material for "Adsorption-driven deformation and landing-footprints of the RBD proteins in SARS-CoV-2 variants onto biological and inanimate surfaces"

[o] *These authors share first authorship.*

### Supplementary Tables and Figures

The delta variant (Figure S4) shows a similar behaviour with the WT (Figure S7), however, we remark that 1 more similar residue between the contacts based on the RBD and PBL is found. Namely, for hydrophobic interaction, ASN (169), while for the hydrophilic surface GLY (170).

In respect to the oscillating behaviour of the delta variant (Figure S5) onto the hydrophilic surfaces is substantially larger than the one shown in the WT variant (Figure S8).

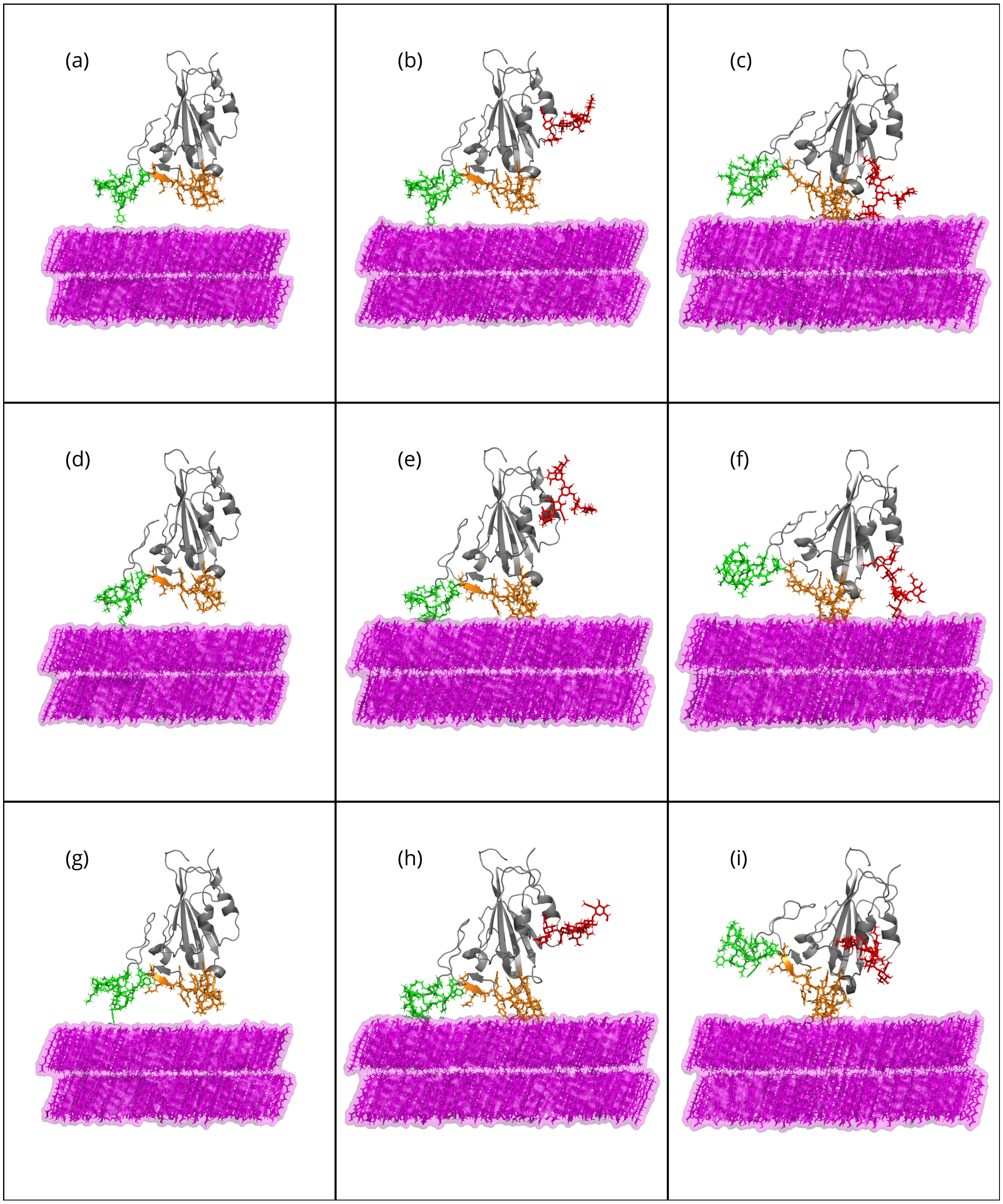


Figure S1: Side view snapshots of the RBD-PBL simulations performed for this research with the hydrophilic (PBL1) substrate at the beginning of the MD production. Rows show snapshots of the RBDs of (a-c) WT,(d-f) Delta, and (g-i) Omicron with the substrate alone, standing vertically to the substrate with its glycan, and rotated respect to the substrate with glycan.

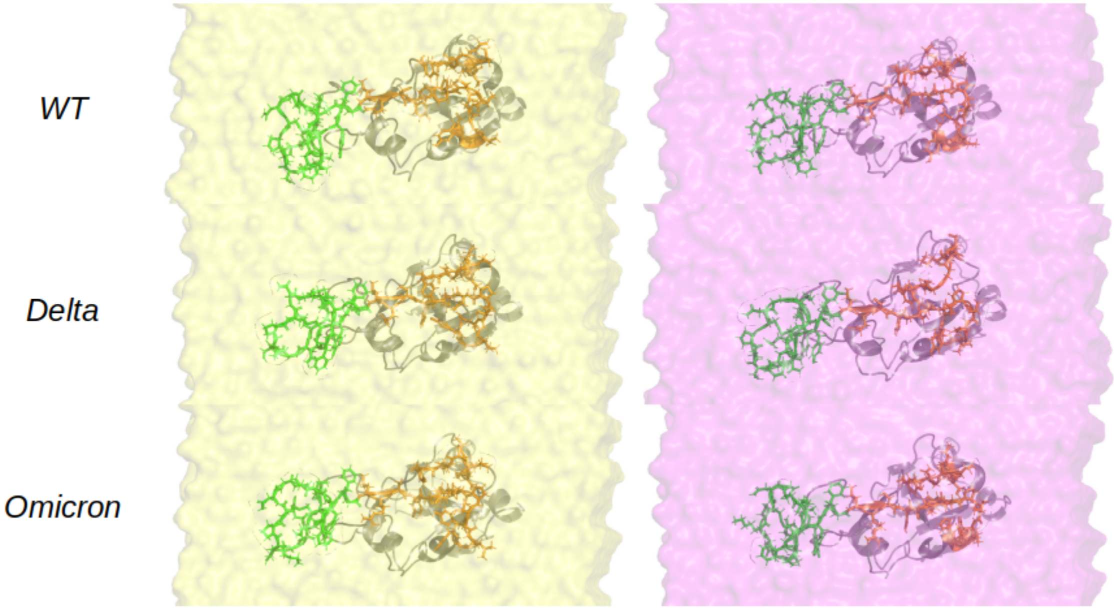


Figure S2: Bottom snapshots of the adsorption of RBDs to the hydrophobic (left) and hydrophilic (right) after 300ns. Ordered as WT, delta, omicron from top to bottom.

Table S1: Table showing the mean contact area, in $nm^2$, over the last 200ns (and their corresponding standard deviation) of the WT, Delta and Omicron-RBDs in presence of the PBL0, PBL1 and ACE2. Data of the simulations in presence of PBL0 and PBL1 with Glycans attached in the RBDs are also added.

| Mean ± Std ($nm^2$) | RBD-PBL0 | RBD-PBL1 | RBD-PBL0 (vGlyc) | RBD-PBL1 (vGlyc) | RBD-ACE2 |
|---|---|---|---|---|---|
| WT | 7 ± 1 | 5 ± 1 | 6 ± 1 | 4 ± 2 | 9 ± 1 |
| Delta | 7 ± 1 | 5 ± 2 | 6 ± 1 | 5 ± 2 | 9 ± 1 |
| Omicron | 8 ± 1 | 5 ± 2 | 7 ± 1 | 6 ± 1 | 9.8 ± 0.5 |

Table S2: Table showing the mean distance (and standard deviation), in Å, of the group regions 1 and 2 during the adsorption over the last 200 ns for the WT, Delta, and Omicron RBDs in presence of PBL0, and PBL1. Adsorption to the surface are considered only in distances lower than 15 Å.

| | Group 1 | Group 2 |
|---|---|---|
| WT-PBL0 | 7.7 ± 0.2 | 9.1 ± 0.2 |
| Delta-PBL0 | 8.0 ± 0.3 | 9.2 ± 0.2 |
| Omicron-PBL0 | 8.1 ± 0.3 | 8.9 ± 0.2 |
| WT-PBL1 | 9.5 ± 0.5 | 9.6 ± 0.5 |
| Delta-PBL1 | 10 ± 1 | 10 ± 1 |
| Omicron-PBL1 | 9.7 ± 0.6 | 10.0 ± 0.6 |

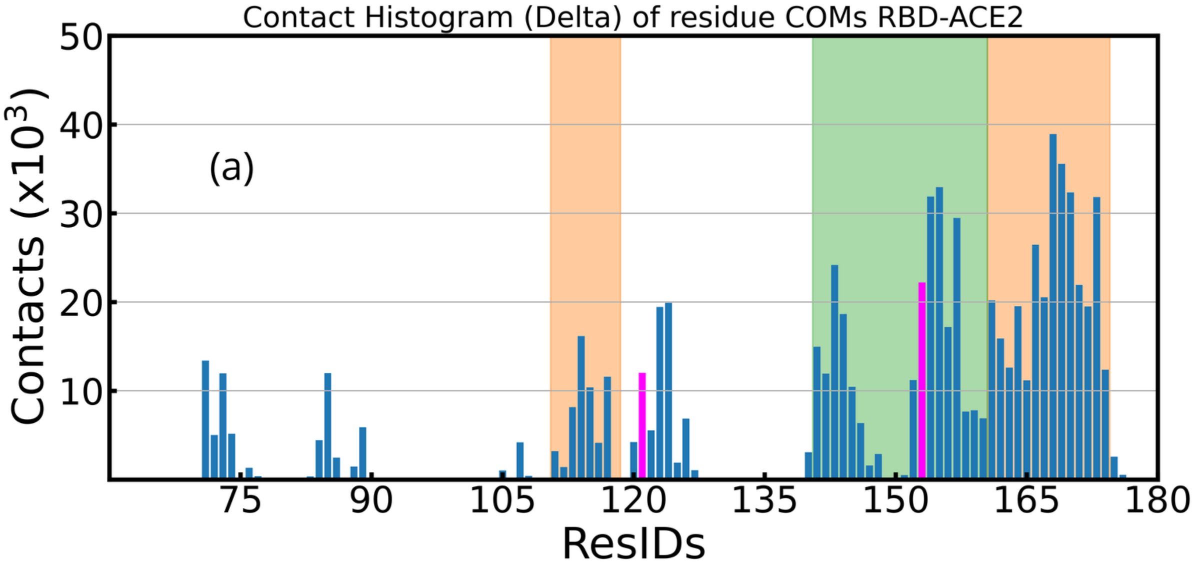


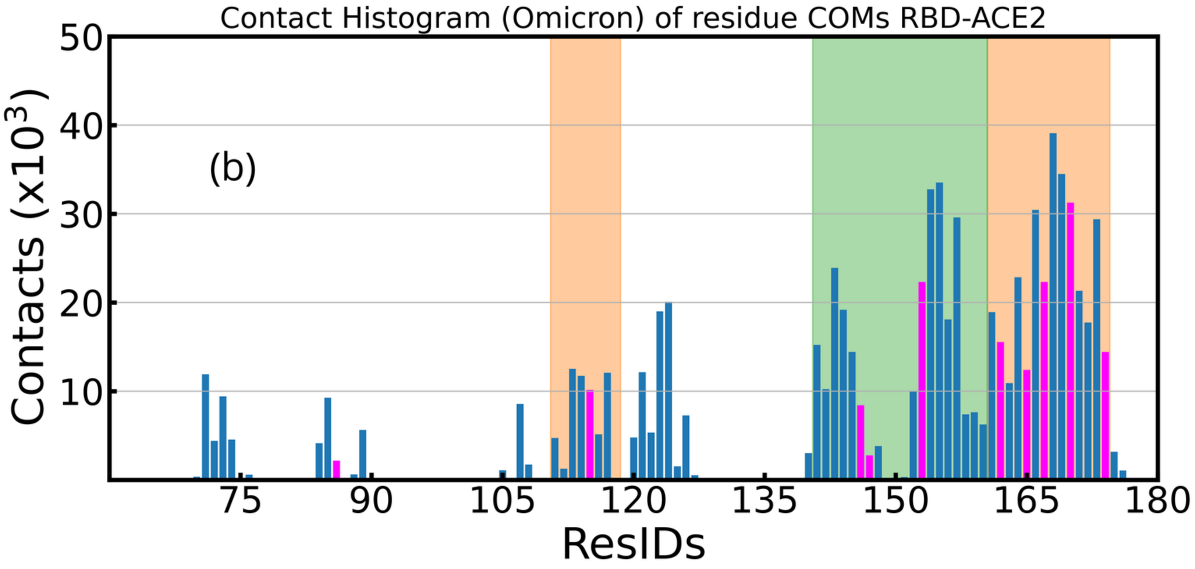


Figure S3: The histograms of contacts between the RBDs (a) delta, and (b) omicron RBDs and the ACE2. Note that the residue mutations to the WT variant are colored in fuchsia.

The Glycan Reader sequance (GRS) of the used Glycans is:

1 BGLCNA

2 - 16A:AFUC

3 - 14B:BGLCNA

4 - - 14B:BMAN

5 - - - 16A:AMAN

6 - - - -12B:BGLCNA

7 - - - 13A:AMAN

8 - - - - 12B:BGLCNA

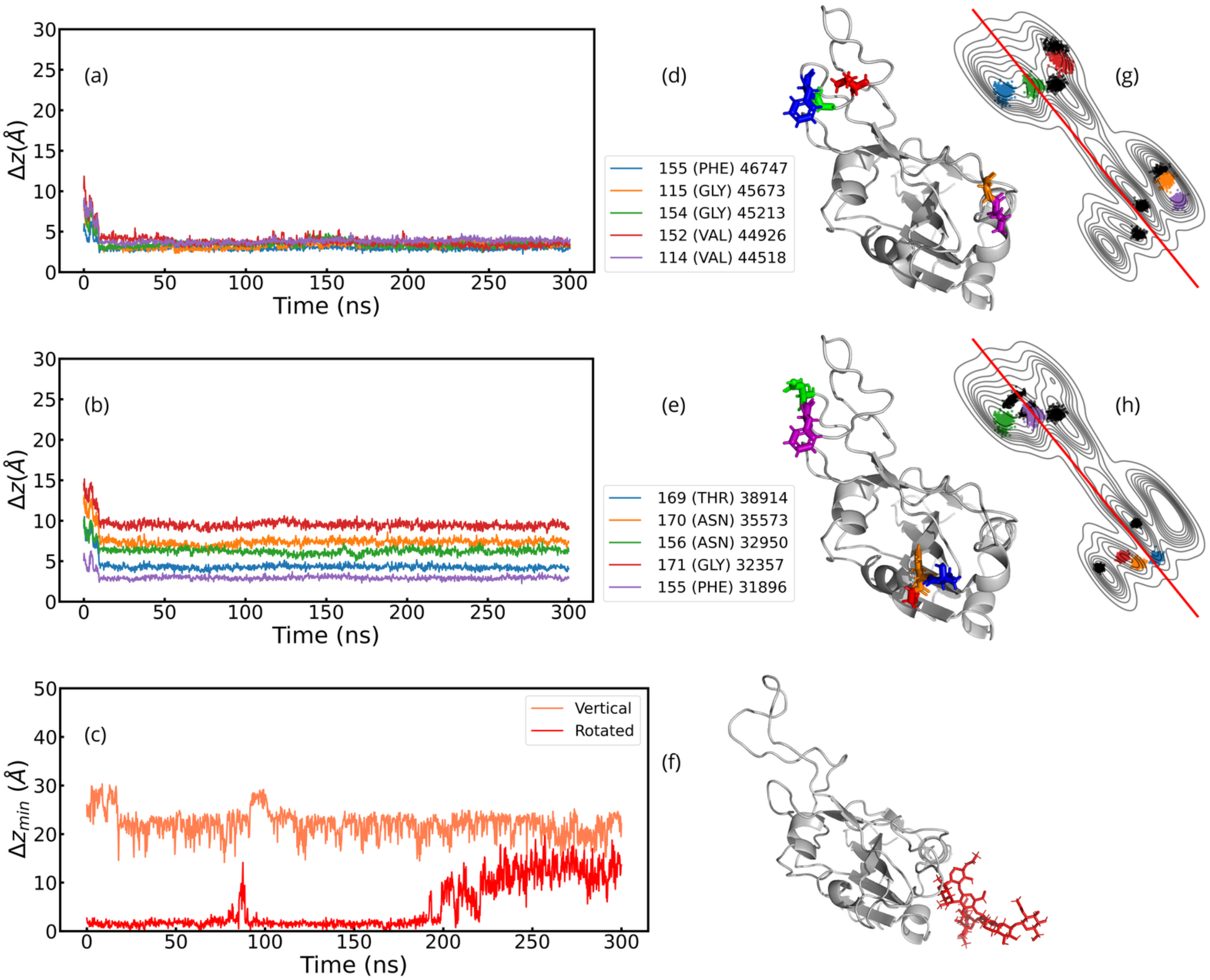


Figure S4: (a,b) Center of mass distance of the residues of Delta-RBD to the hydrophobic substrate of the top 5 residues with most contacts with (a) the hydrophobic substrate and (b) the ACE2. Legends show the ResIDs, the residue names and the total contacts over the trajectory of each ranked residue (format: ResID-ResName TotalContacts). Visualization of residues loci are also shown in (d) and (e) with colors corresponding with the distance plots (a-b). (c) shows the minimum distance between the glycan and the substrate in the hydrophobic surface in a vertical and rotated configuration. In (f), the glycan is shown in red. Note that all snapshots in this plot were taken from a bottom perspective.

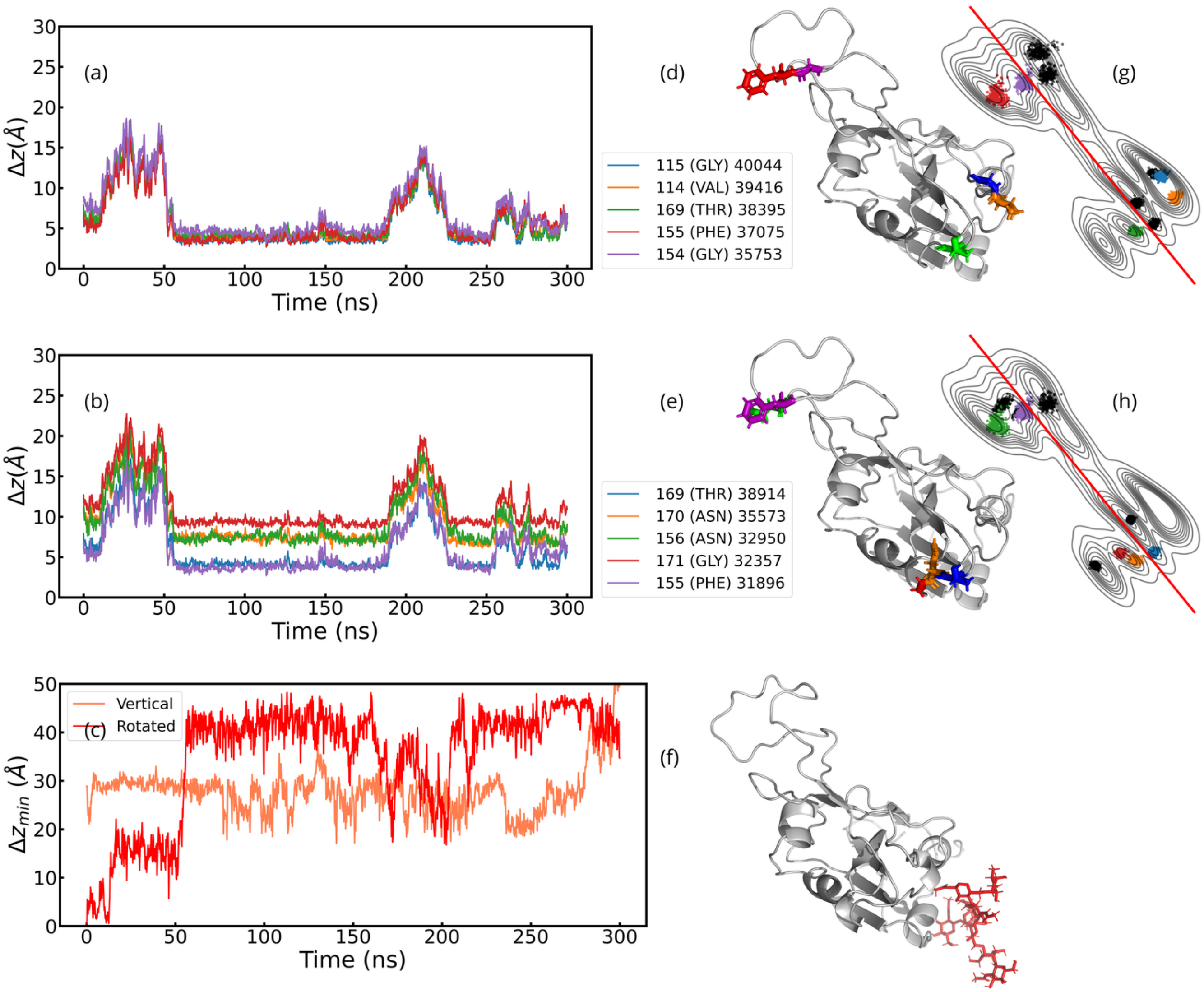


Figure S5: (a,b) Center of mass distance of the residues of Delta-RBD to the hydrophilic substrate of the top 5 residues with most contacts with (a) the hydrophilic substrate and (b) the ACE2. Legends show the ResIDs, the residue names and the total contacts over the trajectory of each ranked residue (format: ResID-ResName TotalContacts). Visualization of residues loci are also shown in (d) and (e) with colors corresponding with the distance plots (a-b). (c) shows the minimum distance between the glycan and the substrate in the hydrophilic surface in a vertical and rotated configuration. In (f), the glycan is shown in red. Note that all snapshots in this plot were taken from a bottom perspective.

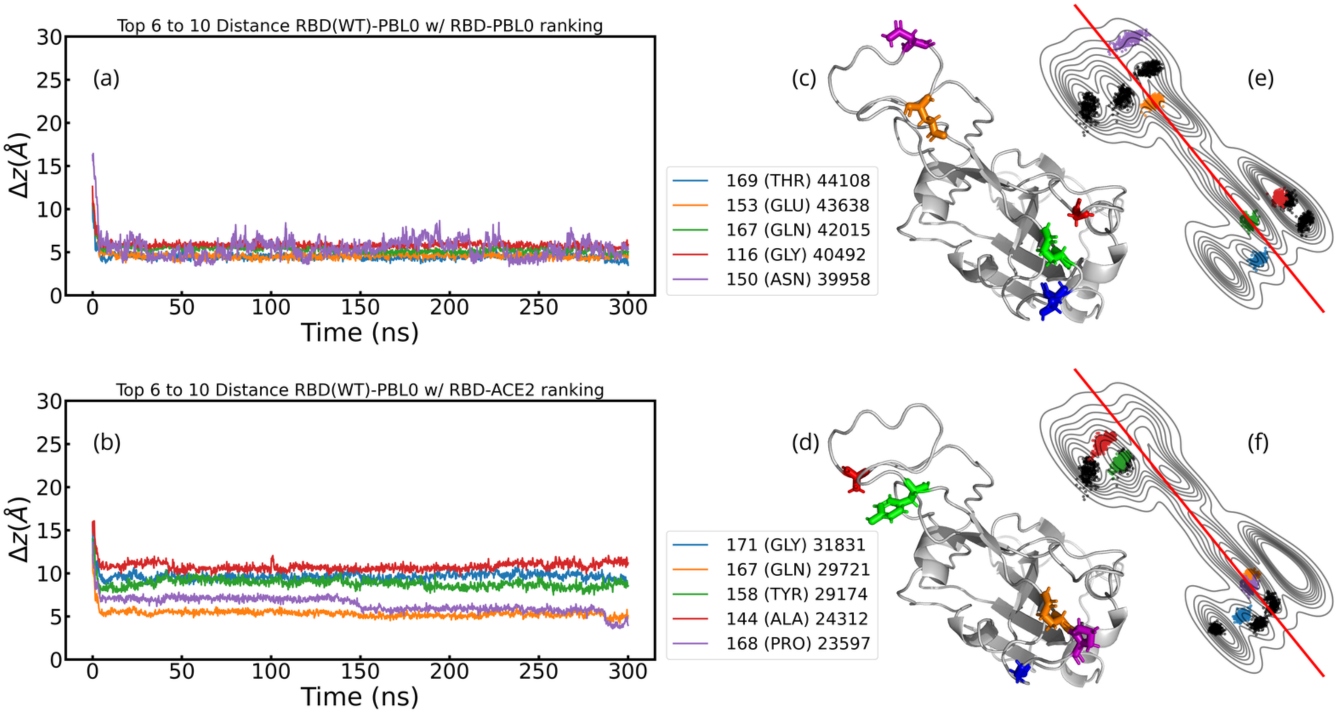


Figure S6: (a,b) Center of mass distance of the residues of WT-RBD to the hydrophobic substrate of the top 6 to 10 residues with most contacts with (a) the hydrophobic substrate and (b) the ACE2. Legends show the ResIDs, the residue names and the total contacts over the trajectory of each ranked residue (format: ResID-ResName TotalContacts). Visualization of residues loci are also shown in (c) and (d) with colors corresponding with the distance plots (a-b).

Table S3: Table contains Normalization factor $N_{\mathrm{var}}/N_{\mathrm{maxvar}}$ for distances 6 Å, 10 Å, 14 Å from PBL0, where $N_{\mathrm{var}}$ and $N_{\mathrm{maxvar}}$ are the number of residues for each variant within the distance ranges and the number of residues within the distance ranges of the variant with maximum residues.

| | Dist. 6 Å | Dist. 10 Å | Dist. 14 Å |
|---|---|---|---|
| WT | 1.00 | 1.00 | 1.00 |
| Delta | 0.92 | 1.00 | 0.98 |
| Omicron | 1.00 | 0.93 | 1.00 |

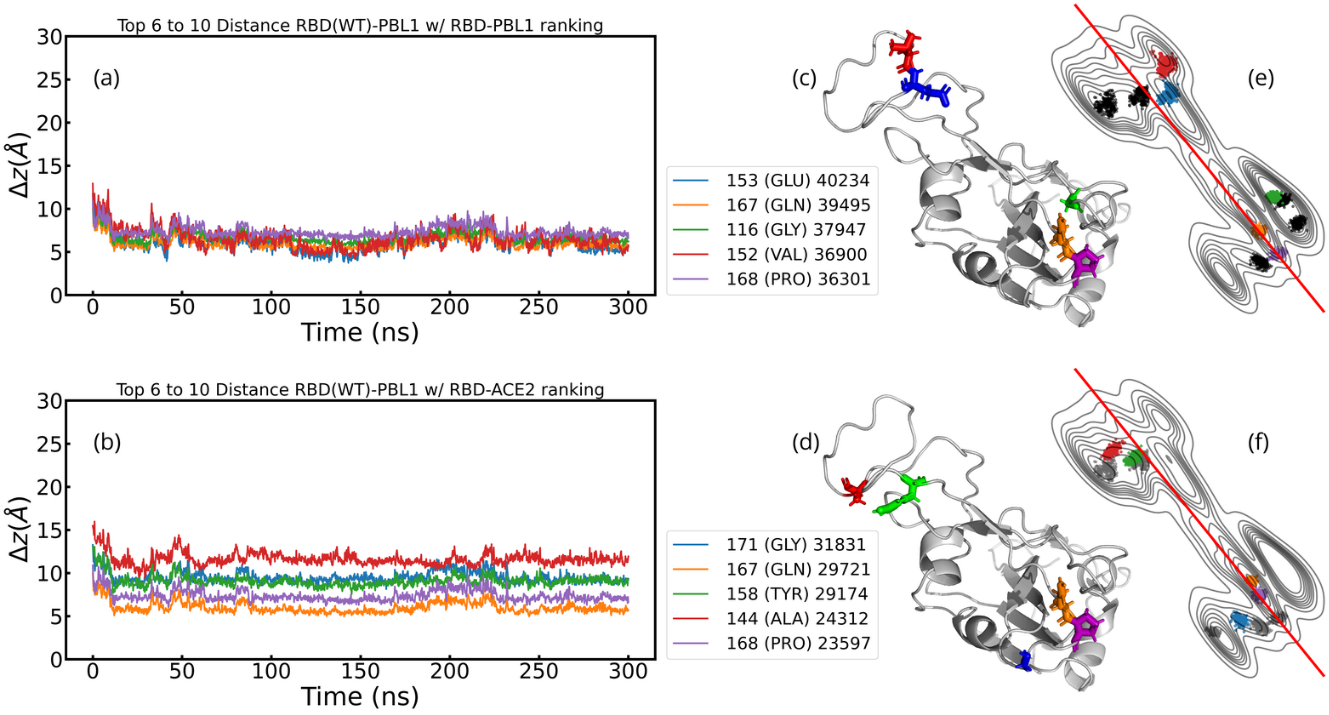


Figure S7: (a,b) Center of mass distance of the residues of WT-RBD to the hydrophilic substrate of the top 6 to 10 residues with most contacts with (a) the hydrophilic substrate and (b) the ACE2. Legends show the ResIDs, the residue names and the total contacts over the trajectory of each ranked residue (format: ResID-ResName TotalContacts).Visualization of residues loci are also shown in (c) and (d) with colors corresponding with the distance plots (a-b).

Table S4: Table contains Normalization factor $N_{\mathrm{var}}/N_{\mathrm{maxvar}}$ for distances 6 Å, 10 Å, 14 Å from PBL1, where $N_{\mathrm{var}}$ and $N_{\mathrm{maxvar}}$ are the number of residues for each variant within the distance ranges and the number of residues within the distance ranges of the variant with maximum residues.

| | Dist. 6 Å | Dist. 10 Å | Dist. 14 Å |
|---|---|---|---|
| WT | 1.00 | 1.00 | 1.00 |
| Delta | 0.57 | 0.82 | 1.00 |
| Omicron | 0.71 | 0.91 | 0.90 |

Table S5: Table contains normalized ratio values for PBL0, using normalization factors on Table S3.

| | Dist. 6 Å | Dist. 10 Å | Dist. 14 Å |
|---|---|---|---|
| WT | 1.50 | 1.26 | 1.13 |
| Delta | 1.42 | 1.15 | 1.06 |
| Omicron | 1.50 | 1.19 | 1.09 |

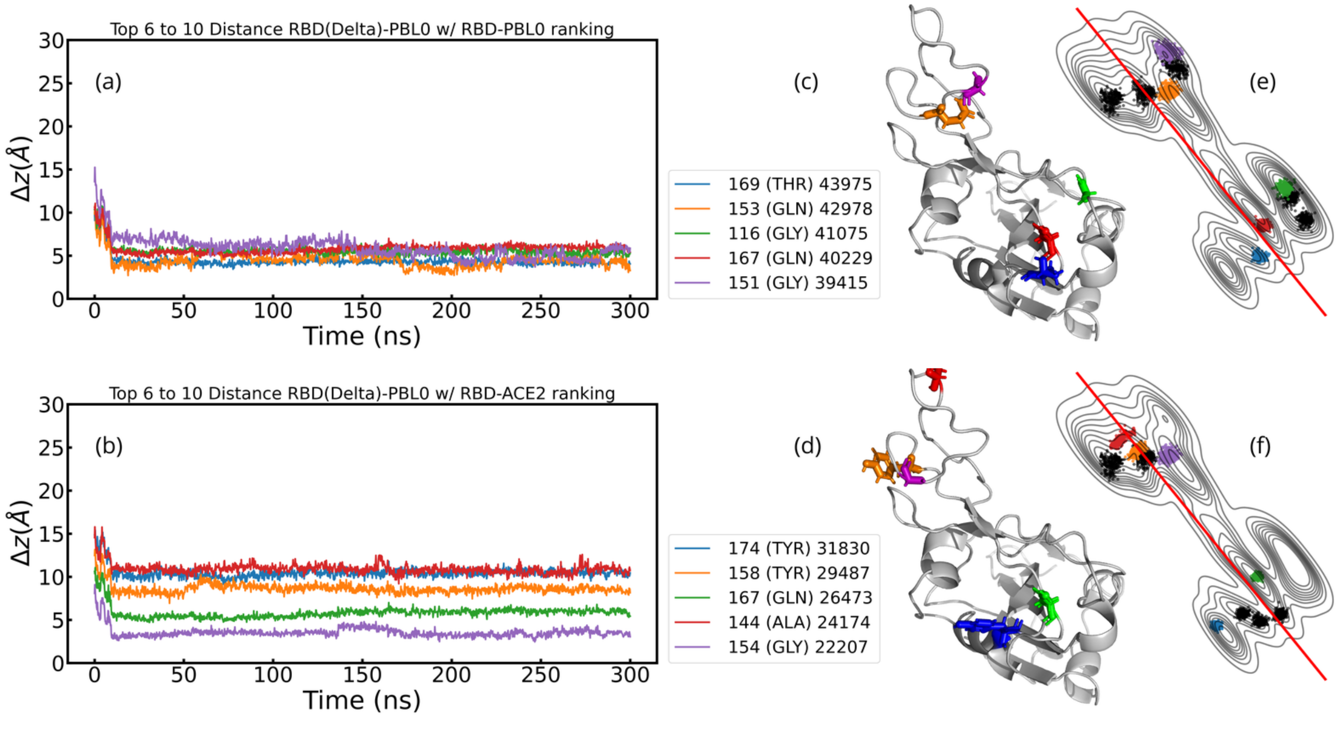


Figure S8: (a,b) Center of mass distance of the residues of Delta-RBD to the hydrophobic substrate of the top 6 to 10 residues with most contacts with (a) the hydrophobic substrate and (b) the ACE2. Legends show the ResIDs, the residue names and the total contacts over the trajectory of each ranked residue (format: ResID-ResName TotalContacts). Visualization of residues loci are also shown in (c) and (d) with colors corresponding with the distance plots (a-b).

Table S6: Table contains normalized ratio values for PBL1, using normalization factors on Table S4.

| | Dist. 6 Å | Dist. 10 Å | Dist. 14 Å |
|---|---|---|---|
| WT | 1.71 | 1.36 | 1.17 |
| Delta | 1.86 | 1.27 | 1.14 |
| Omicron | 1.71 | 1.32 | 1.12 |

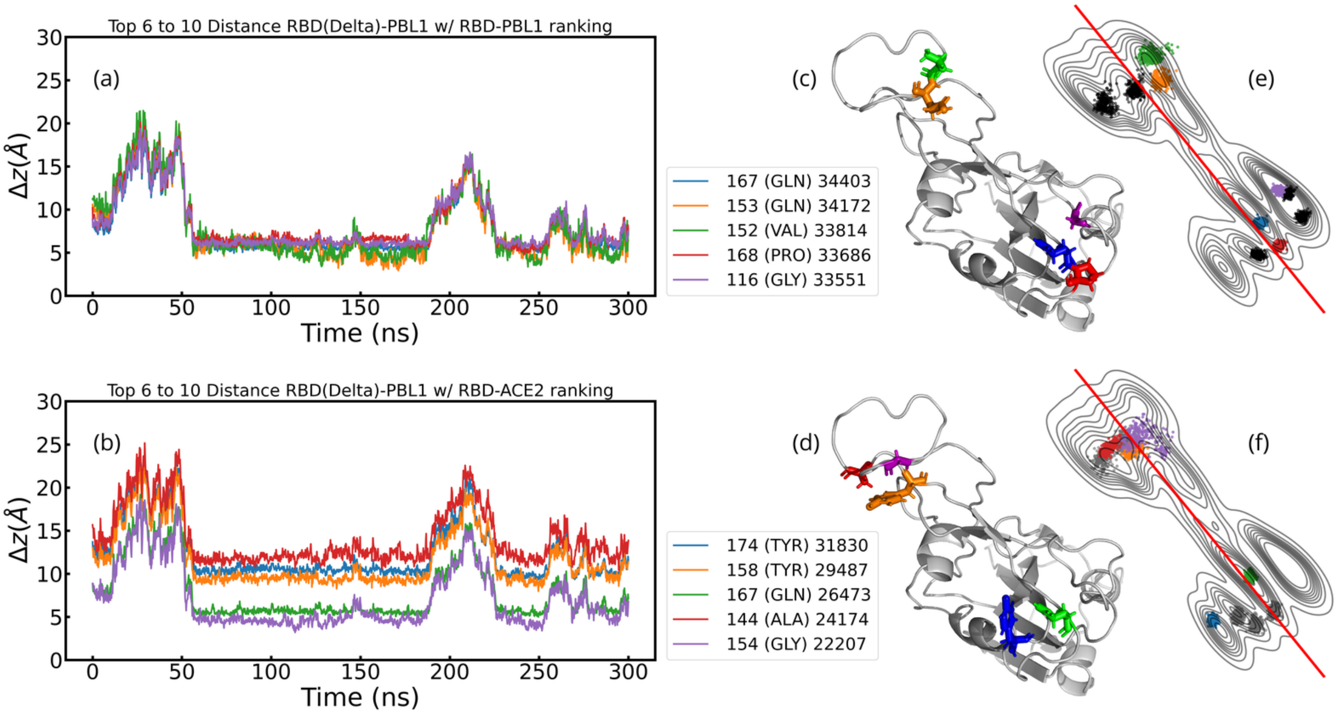


Figure S9: (a,b) Center of mass distance of the residues of Delta-RBD to the hydrophilic substrate of the top 6 to 10 residues with most contacts with (a) the hydrophilic substrate and (b) the ACE2. Legends show the ResIDs, the residue names and the total contacts over the trajectory of each ranked residue (format: ResID-ResName TotalContacts). Visualization of residues loci are also shown in (c) and (d) with colors corresponding with the distance plots (a-b).

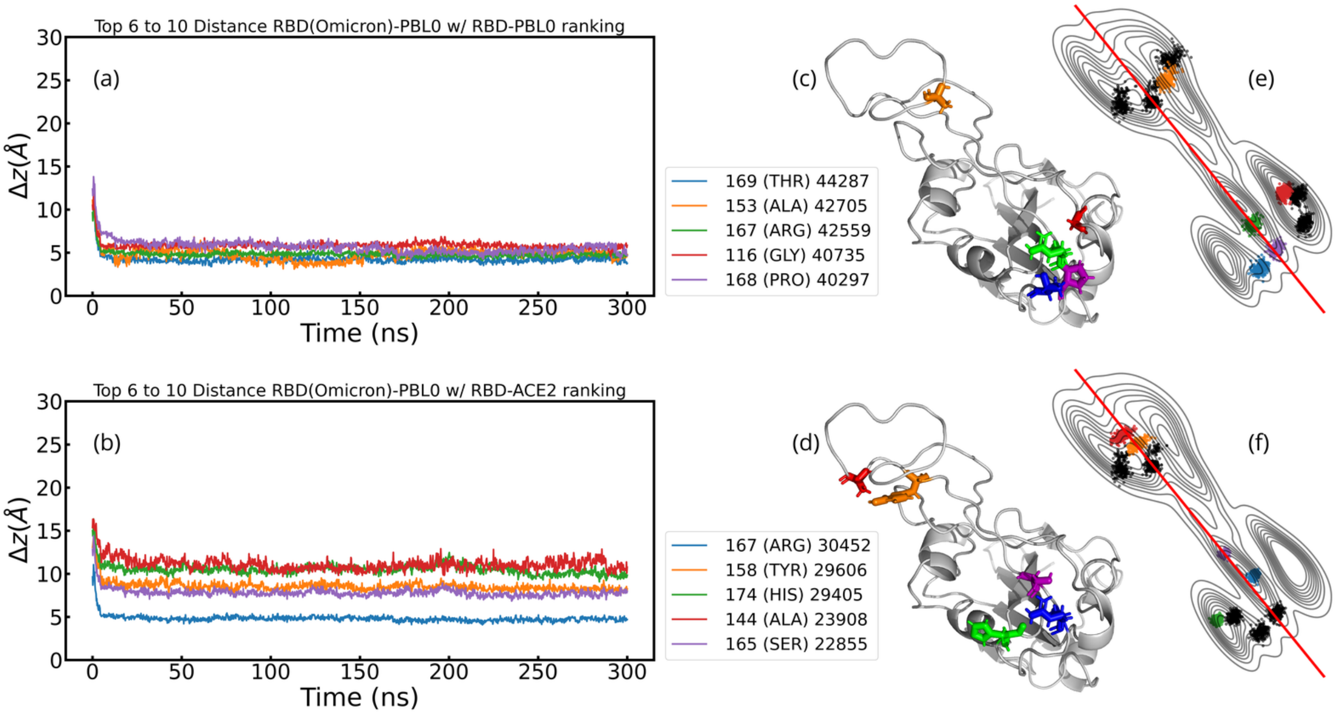


Figure S10: (a,b) Center of mass distance of the residues of Omicron-RBD to the hydrophobic substrate of the top 6 to 10 residues with most contacts with (a) the hydrophobic substrate and (b) the ACE2. Legends show the ResIDs, the residue names and the total contacts over the trajectory of each ranked residue (format: ResID-ResName TotalContacts). Visualization of residues loci are also shown in (c) and (d) with colors corresponding with the distance plots (a-b).

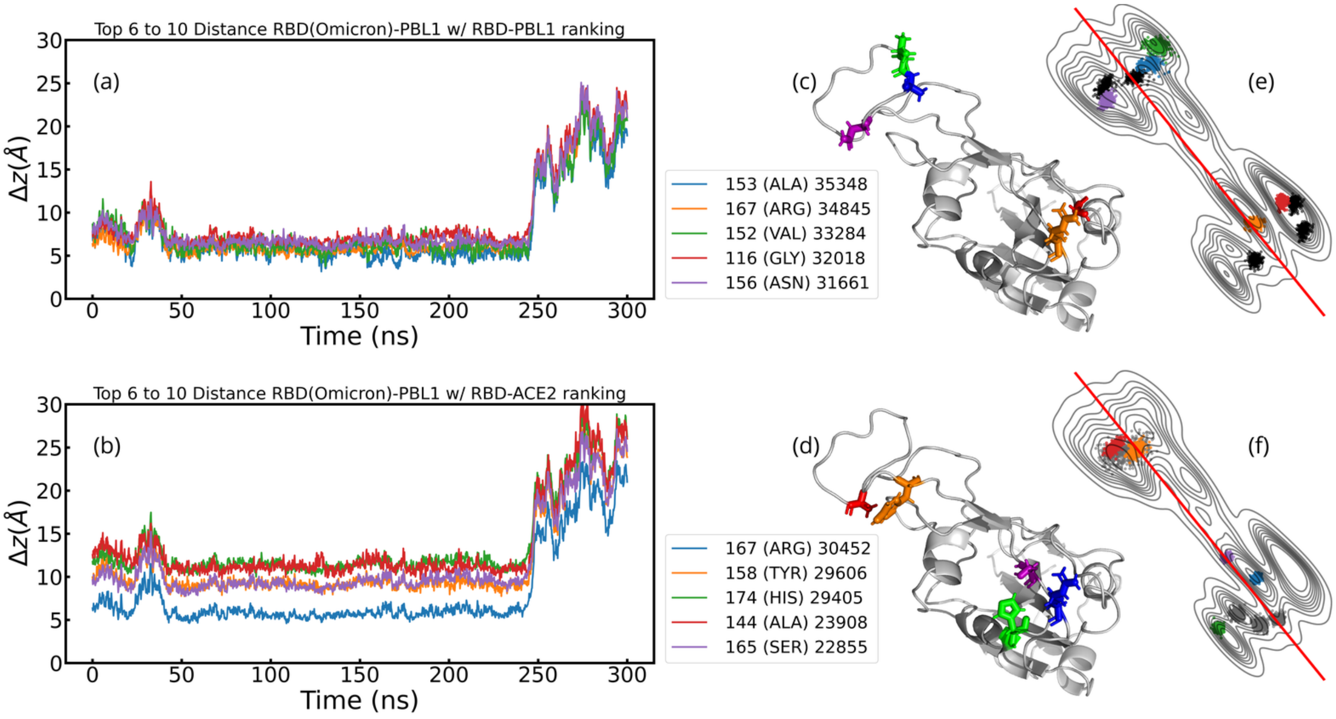


Figure S11: (a,b) Center of mass distance of the residues of Omicron-RBD to the hydrophilic substrate of the top 6 to 10 residues with most contacts with (a) the hydrophilic substrate and (b) the ACE2. Legends show the ResIDs, the residue names and the total contacts over the trajectory of each ranked residue (format: ResID-ResName TotalContacts). Visualization of residues loci are also shown in (c) and (d) with colors corresponding with the distance plots (a-b).

Table S7: Table contains the ResIDs and corresponding residue names of residues used in Group 1. Mutations are highlighted in fuchsia. **CN**: Charged Negative, **CP**: Charged Positive, **UP**: Uncharged Polar and **NP**: NonPolar

| ResIDs | WT resnames | Delta resnames | Omicron resnames |
|---|---|---|---|
| 141 | ILE (NP) | ILE (NP) | ILE (NP) |
| 142 | TYR (UP) | TYR (UP) | TYR (UP) |
| 143 | GLN (UP) | GLN (UP) | GLN (UP) |
| 144 | ALA (NP) | ALA (NP) | ALA (NP) |
| 145 | GLY (NP) | GLY (NP) | GLY (NP) |
| 146 | SER (UP) | SER (UP) | ASN (UP) |
| 147 | THR (UP) | THR (UP) | LYS (CP) |
| 148 | PRO (NP) | PRO (NP) | PRO (NP) |
| 149 | CYS (NP) | CYS (NP) | CYS (NP) |
| 150 | ASN (UP) | ASN (UP) | ASN (UP) |
| 151 | GLY (NP) | GLY (NP) | GLY (NP) |
| 152 | VAL (NP) | VAL (NP) | VAL (NP) |
| 153 | GLU (CN) | GLN (UP) | ALA (NP) |
| 154 | GLY (NP) | GLY (NP) | GLY (NP) |
| 155 | PHE (NP) | PHE (NP) | PHE (NP) |
| 156 | ASN (UP) | ASN (UP) | ASN (UP) |
| 157 | CYS (NP) | CYS (NP) | CYS (NP) |
| 158 | TYR (UP) | TYR (UP) | TYR (UP) |
| 159 | PHE (NP) | PHE (NP) | PHE (NP) |
| 160 | PRO (NP) | PRO (NP) | PRO (NP) |

Table S8: Table contains the ResIDs and corresponding residue names of residues used in Group 2. Mutations are highlighted in fuchsia. **CN**: Charged Negative, **CP**: Charged Positive, **UP**: Uncharged Polar and **NP**: NonPolar

| ResIDs | WT resnames | Delta resnames | Omicron resnames |
|---|---|---|---|
| 111 | ASP (CN) | ASP (CN) | ASP (CN) |
| 112 | SER (UP) | SER (UP) | SER (UP) |
| 113 | LYS (CP) | LYS (CP) | LYS (CP) |
| 114 | VAL (NP) | VAL (NP) | VAL (NP) |
| 115 | GLY (NP) | GLY (NP) | SER (UP) |
| 116 | GLY (NP) | GLY (NP) | GLY (NP) |
| 118 | TYR (UP) | TYR (UP) | TYR (UP) |
| 117 | ASN (UP) | ASN (UP) | ASN (UP) |
| 161 | LEU (NP) | LEU (NP) | LEU (NP) |
| 162 | GLN (UP) | GLN (UP) | ARG (CP) |
| 163 | SER (UP) | SER (UP) | SER (UP) |
| 164 | TYR (UP) | TYR (UP) | TYR (UP) |
| 165 | GLY (NP) | GLY (NP) | SER (UP) |
| 166 | PHE (NP) | PHE (NP) | PHE (NP) |
| 167 | GLN (UP) | GLN (UP) | ARG (CP) |
| 168 | PRO (NP) | PRO (NP) | PRO (NP) |
| 169 | THR (UP) | THR (UP) | THR (UP) |
| 170 | ASN (UP) | ASN (UP) | TYR (UP) |
| 171 | GLY (NP) | GLY (NP) | GLY (NP) |
| 172 | VAL (NP) | VAL (NP) | VAL (NP) |
| 173 | GLY (NP) | GLY (NP) | GLY (NP) |
| 174 | TYR (UP) | TYR (UP) | HIS (CP) |

Table S9: Table contains the ResIDs and corresponding residue names and type of residues that define the Crook-handle for PBL0. Mutations are highlighted in fuchsia. Note that for omicron, this region is a hydrophobic pocket. Residue types are given by **CN**: Charged Negative, **CP**: Charged Positive, **UP**: Uncharged Polar and **NP**: NonPolar.

| Polarity 0 | | | |
|---|---|---|---|
| ResIDs | WT resnames | Delta resnames | Omicron resnames |
| 150 | ASN (UP) | ASN (UP) | ASN (UP) |
| 151 | GLY (NP) | GLY (NP) | GLY (NP) |
| 152 | VAL (NP) | VAL (NP) | VAL (NP) |
| 153 | GLU (CN) | GLN (UP) | ALA (NP) |
| 154 | GLY (NP) | GLY (NP) | GLY (NP) |
| 155 | PHE (NP) | PHE (NP) | PHE (NP) |

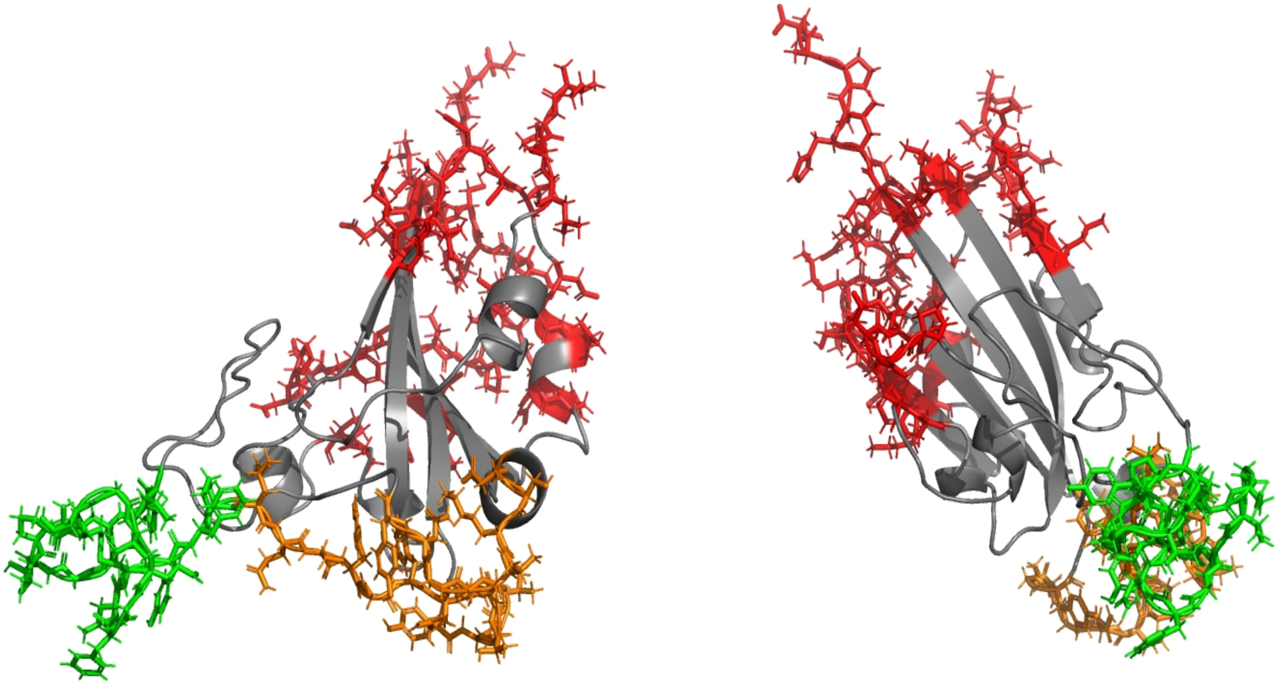

Figure S12: Front (left) and side view (right) of the RBD showing Group 1, Group 2 and restraint residues in green, orange and red, respectively.

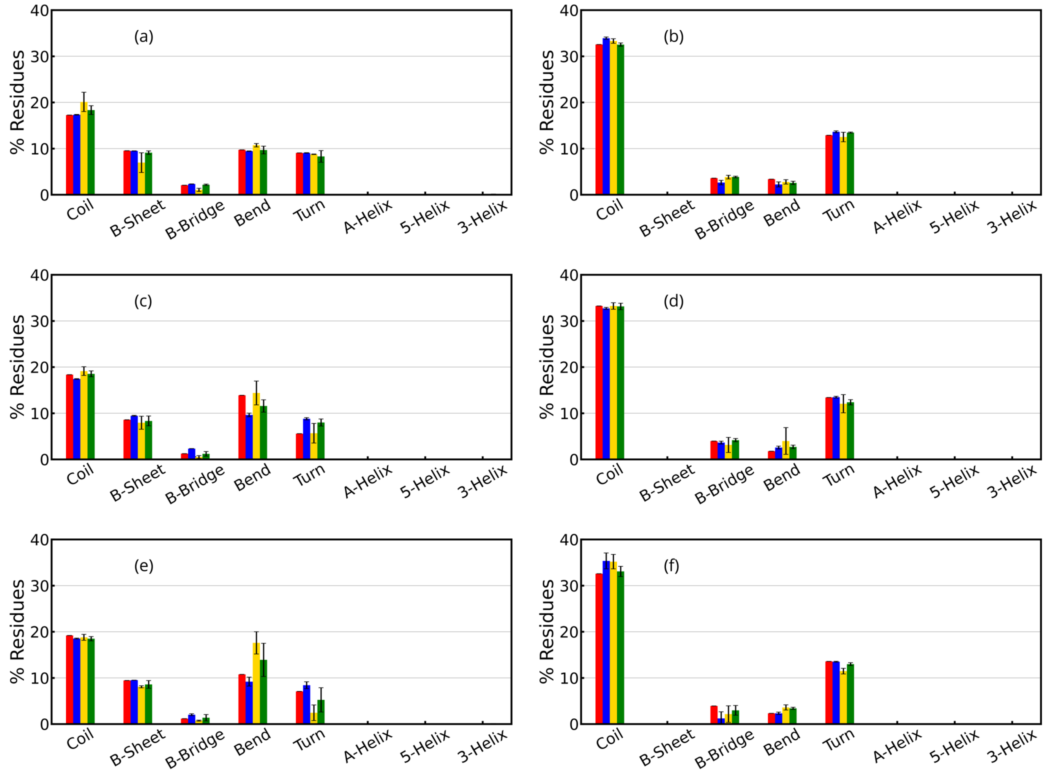


Figure S13: Second Structure percentage (SS %) of group 1 (left column) and group 2 (right column) averaged over the last 200 ns of the simulations (with the exception of the simulations of RBD alone, is the mean of all its 20 ns long trajectory) for (a,b) WT, (c,d) Delta, and (e,f) Omicron variants alone in water (red), with the ACE2 (blue), in presence of PBL0 (yellow), and PBL1 (green). Values are normalized over the number of residues in group 1 and 2.

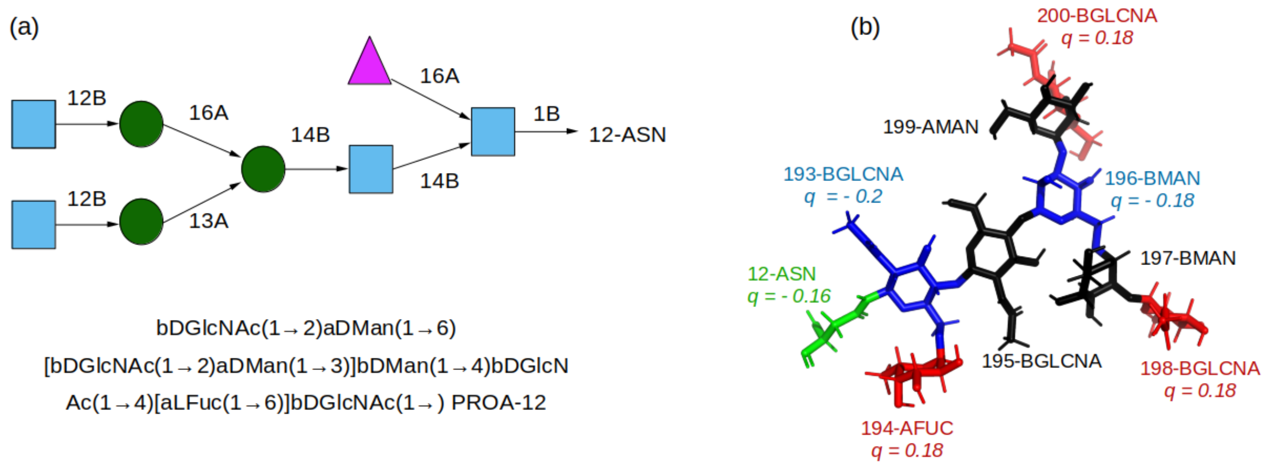


Figure S14: (a) Glycan Structure. (b) Glycan mapping and charge distribution. In (b), green is residue 12-ASN that binds the glycan to the protein, and the glycan is colored according the charge type. Red are positively charged, blue negatively charged and black are neutrally charged carbohydrates.

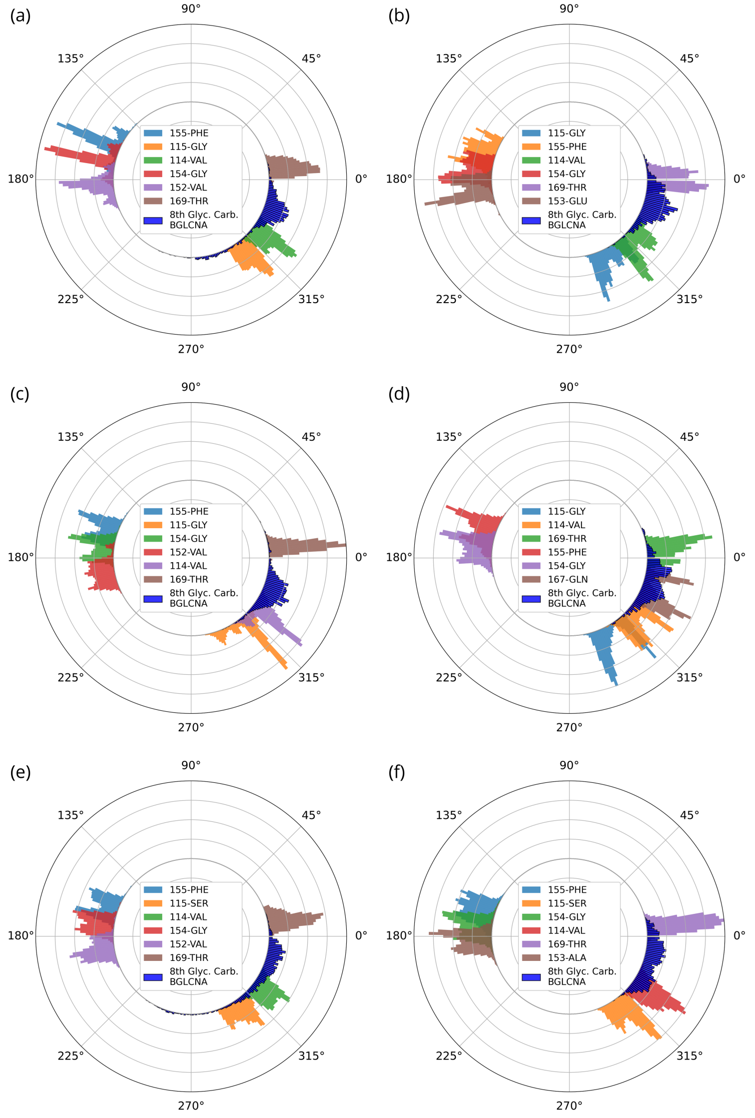


Figure S15: Angular Histogram of top contact residues of (a,b) WT, (c,d) Delta, and (e,f) Omicron. Center point is set at 20 Å distance from the mean position over time of each selected residue COM of the RBDs for each PBL (left column for hydrophobic surface and right for hydrophilic surfaces), and the glycan histogram from RBD-PBLs with Glycans simulations. Residues selected are Top 6 residues with most contacts to the PBLs, and the 8th residue of the glycan which is one of the further residues from the binding residue 12-ASN Schematic of the Glycan can be visualized in SI (Figure S14). Note that the first five colors are consistent to residue colors in Figure 11. Histogram is normalized over the maximum value for each surface, i.e. the maximum value of 169-THR in Delta-PBL0 and Omicron-PBL1.

Table S10: Table listing the ResIDs and corresponding residue names for each variant. Left and right legs are highlighted in green and orange rows, respectively. Mutations of delta and omicron referenced to wild-type of the RBD are shown with fuchsia highlighted cells. Fourth column indicates if residue is restraint in the model. It must be noted that ResIDs of the RBDs were initialized to one for practical reasons. Analogue values to PDB ID:6VSB requires adding up 331 units to the ResIDs,i.e. 1-ILE → 332-ILE (PDB) and 192-THR→ 523-THR.

| ResIDs | WT | Delta | Omicron | Restraint |
|---|---|---|---|---|
| 1 | ILE | ILE | ILE | Yes |
| 2 | THR | THR | THR | Yes |
| 3 | ASN | ASN | ASN | Yes |
| 4 | LEU | LEU | LEU | Yes |
| 5 | CYS | CYS | CYS | No |
| 6 | PRO | PRO | PRO | No |
| 7 | PHE | PHE | PHE | No |
| 8 | GLY | GLY | ASP | No |
| 9 | GLU | GLU | GLU | No |
| 10 | VAL | VAL | VAL | No |
| 11 | PHE | PHE | PHE | No |
| 12 | ASN | ASN | ASN | No |
| 13 | ALA | ALA | ALA | No |
| 14 | THR | THR | THR | No |
| 15 | ARG | ARG | ARG | No |
| 16 | PHE | PHE | PHE | No |
| 17 | ALA | ALA | ALA | No |
| 18 | SER | SER | SER | No |
| 19 | VAL | VAL | VAL | No |
| 20 | TYR | TYR | TYR | No |
| 21 | ALA | ALA | ALA | No |
| 22 | TRP | TRP | TRP | No |
| 23 | ASN | ASN | ASN | No |
| 24 | ARG | ARG | ARG | No |
| 25 | LYS | LYS | LYS | Yes |
| 26 | ARG | ARG | ARG | Yes |
| 27 | ILE | ILE | ILE | Yes |
| 28 | SER | SER | SER | Yes |
| 29 | ASN | ASN | ASN | Yes |
| 30 | CYS | CYS | CYS | Yes |
| 31 | VAL | VAL | VAL | Yes |
| 32 | ALA | ALA | ALA | No |
| 33 | ASP | ASP | ASP | Yes |
| 34 | TYR | TYR | TYR | Yes |
| 35 | SER | SER | SER | Yes |
| 36 | VAL | VAL | VAL | Yes |

| ResIDs | WT | Delta | Omicron | Restraint |
|---|---|---|---|---|
| 37 | LEU | LEU | LEU | No |
| 38 | TYR | TYR | TYR | Yes |
| 39 | ASN | ASN | ASN | Yes |
| 40 | SER | SER | LEU | No |
| 41 | ALA | ALA | ALA | No |
| 42 | SER | SER | PRO | No |
| 43 | PHE | PHE | PHE | No |
| 44 | SER | SER | PHE | No |
| 45 | THR | THR | THR | No |
| 46 | PHE | PHE | PHE | No |
| 47 | LYS | LYS | LYS | Yes |
| 48 | CYS | CYS | CYS | Yes |
| 49 | TYR | TYR | TYR | Yes |
| 50 | GLY | GLY | GLY | Yes |
| 51 | VAL | VAL | VAL | Yes |
| 52 | SER | SER | SER | Yes |
| 53 | PRO | PRO | PRO | Yes |
| 54 | THR | THR | THR | Yes |
| 55 | LYS | LYS | LYS | Yes |
| 56 | LEU | LEU | LEU | Yes |
| 57 | ASN | ASN | ASN | Yes |
| 58 | ASP | ASP | ASP | Yes |
| 59 | LEU | LEU | LEU | Yes |
| 60 | CYS | CYS | CYS | Yes |
| 61 | PHE | PHE | PHE | Yes |
| 62 | THR | THR | THR | Yes |
| 63 | ASN | ASN | ASN | Yes |
| 64 | VAL | VAL | VAL | No |
| 65 | TYR | TYR | TYR | No |
| 66 | ALA | ALA | ALA | No |
| 67 | ASP | ASP | ASP | No |
| 68 | SER | SER | SER | No |
| 69 | PHE | PHE | PHE | No |
| 70 | VAL | VAL | VAL | No |
| 71 | ILE | ILE | ILE | No |
| 72 | ARG | ARG | ARG | No |
| 73 | GLY | GLY | GLY | No |
| 74 | ASP | ASP | ASP | No |
| 75 | GLU | GLU | GLU | No |
| 76 | VAL | VAL | VAL | No |
| 77 | ARG | ARG | ARG | No |
| 78 | GLN | GLN | GLN | No |
| 79 | ILE | ILE | ILE | No |

| ResIDs | WT | Delta | Omicron | Restraint |
|---|---|---|---|---|
| 80 | ALA | ALA | ALA | Yes |
| 81 | PRO | PRO | PRO | Yes |
| 82 | GLY | GLY | GLY | No |
| 83 | GLN | GLN | GLN | No |
| 84 | THR | THR | THR | No |
| 85 | GLY | GLY | GLY | No |
| 86 | LYS | LYS | ASN | No |
| 87 | ILE | ILE | ILE | No |
| 88 | ALA | ALA | ALA | No |
| 89 | ASP | ASP | ASP | No |
| 90 | TYR | TYR | TYR | No |
| 91 | ASN | ASN | ASN | No |
| 92 | TYR | TYR | TYR | No |
| 93 | LYS | LYS | LYS | No |
| 94 | LEU | LEU | LEU | No |
| 95 | PRO | PRO | PRO | Yes |
| 96 | ASP | ASP | ASP | Yes |
| 97 | ASP | ASP | ASP | Yes |
| 98 | PHE | PHE | PHE | Yes |
| 99 | THR | THR | THR | Yes |
| 100 | GLY | GLY | GLY | No |
| 101 | CYS | CYS | CYS | No |
| 102 | VAL | VAL | VAL | No |
| 103 | ILE | ILE | ILE | No |
| 104 | ALA | ALA | ALA | No |
| 105 | TRP | TRP | TRP | No |
| 106 | ASN | ASN | ASN | No |
| 107 | SER | SER | SER | No |
| 108 | ASN | ASN | ASN | No |
| 109 | ASN | ASN | LYS | No |
| 110 | LEU | LEU | LEU | No |
| 111 | ASP | ASP | ASP | No |
| 112 | SER | SER | SER | No |
| 113 | LYS | LYS | LYS | No |
| 114 | VAL | VAL | VAL | No |
| 115 | GLY | GLY | SER | No |
| 116 | GLY | GLY | GLY | No |
| 117 | ASN | ASN | ASN | No |
| 118 | TYR | TYR | TYR | No |
| 119 | ASN | ASN | ASN | No |
| 120 | TYR | TYR | TYR | No |
| 121 | LEU | ARG | LEU | No |
| 122 | TYR | TYR | TYR | No |

| ResIDs | WT | Delta | Omicron | Restraint |
|---|---|---|---|---|
| 123 | ARG | ARG | ARG | No |
| 124 | LEU | LEU | LEU | No |
| 125 | PHE | PHE | PHE | No |
| 126 | ARG | ARG | ARG | No |
| 127 | LYS | LYS | LYS | No |
| 128 | SER | SER | SER | No |
| 129 | ASN | ASN | ASN | No |
| 130 | LEU | LEU | LEU | No |
| 131 | LYS | LYS | LYS | No |
| 132 | PRO | PRO | PRO | No |
| 133 | PHE | PHE | PHE | No |
| 134 | GLU | GLU | GLU | No |
| 135 | ARG | ARG | ARG | No |
| 136 | ASP | ASP | ASP | No |
| 137 | ILE | ILE | ILE | No |
| 138 | SER | SER | SER | No |
| 139 | THR | THR | THR | No |
| 140 | GLU | GLU | GLU | No |
| 141 | ILE | ILE | ILE | No |
| 142 | TYR | TYR | TYR | No |
| 143 | GLN | GLN | GLN | No |
| 144 | ALA | ALA | ALA | No |
| 145 | GLY | GLY | GLY | No |
| 146 | SER | SER | ASN | No |
| 147 | THR | THR | LYS | No |
| 148 | PRO | PRO | PRO | No |
| 149 | CYS | CYS | CYS | No |
| 150 | ASN | ASN | ASN | No |
| 151 | GLY | GLY | GLY | No |
| 152 | VAL | VAL | VAL | No |
| 153 | GLU | GLN | ALA | No |
| 154 | GLY | GLY | GLY | No |
| 155 | PHE | PHE | PHE | No |
| 156 | ASN | ASN | ASN | No |
| 157 | CYS | CYS | CYS | No |
| 158 | TYR | TYR | TYR | No |
| 159 | PHE | PHE | PHE | No |
| 160 | PRO | PRO | PRO | No |
| 161 | LEU | LEU | LEU | No |
| 162 | GLN | GLN | ARG | No |
| 163 | SER | SER | SER | No |
| 164 | TYR | TYR | TYR | No |
| 165 | GLY | GLY | SER | No |

| ResIDs | WT | Delta | Omicron | Restraint |
|---|---|---|---|---|
| 166 | PHE | PHE | PHE | No |
| 167 | GLN | GLN | ARG | No |
| 168 | PRO | PRO | PRO | No |
| 169 | THR | THR | THR | No |
| 170 | ASN | ASN | TYR | No |
| 171 | GLY | GLY | GLY | No |
| 172 | VAL | VAL | VAL | No |
| 173 | GLY | GLY | GLY | No |
| 174 | TYR | TYR | HIS | No |
| 175 | GLN | GLN | GLN | No |
| 176 | PRO | PRO | PRO | No |
| 177 | TYR | TYR | TYR | No |
| 178 | ARG | ARG | ARG | No |
| 179 | VAL | VAL | VAL | No |
| 180 | VAL | VAL | VAL | No |
| 181 | VAL | VAL | VAL | No |
| 182 | LEU | LEU | LEU | No |
| 183 | SER | SER | SER | No |
| 184 | PHE | PHE | PHE | No |
| 185 | GLU | GLU | GLU | Yes |
| 186 | LEU | LEU | LEU | Yes |
| 187 | LEU | LEU | LEU | Yes |
| 188 | HIS | HIS | HIS | Yes |
| 189 | ALA | ALA | ALA | Yes |
| 190 | PRO | PRO | PRO | Yes |
| 191 | ALA | ALA | ALA | Yes |
| 192 | THR | THR | THR | Yes |
